# Environmental conditions drive selection and recovery following disease-induced declines

**DOI:** 10.64898/2026.07.09.737596

**Authors:** Samantha Hoff, Joseph R. Hoyt, Alex T. Grimaudo, Macy J. Kailing, Nichole A. Laggan, Chris D. Kailing, Allen Kurta, John E. DePue, Alyssa B. Bennett, Heather M. Kaarakka, Jennifer A. Redell, J. Paul White, Ashley R. Meyer, Kate E. Langwig

## Abstract

Emerging infectious diseases threaten public health and biodiversity across the globe^1,2^. Disease outcomes are frequently dependent on local environmental conditions^3–5^, but how these factors shape host adaptation and long-term recovery are often unknown^6^. Here we combine two decades of population, disease, and environmental data with a common garden experiment to investigate the drivers of variable declines and recovery for remnant bat populations following the emergence of the fungal disease, white-nose syndrome. We find that initial declines were greater and faster in warmer sites (88.3% vs. 74.2% in cold sites), but these populations recovered more quickly and hosts developed higher resistance (1.5x reduction of fungal loads) than populations from colder sites that were buffered from initial impacts. Our experimental data suggest that warm sites served as hotspots of host adaptation where selective pressures were stronger because thermal conditions approached optimal growth for the pathogen, which eventually favored the development of high pathogen resistance. Populations in colder sites experienced weaker selective pressure and thus remain more susceptible, although bats from larger colonies were more likely to survive, suggesting that adaptive traits exist in these populations, but at much lower frequency. These findings show that the environmental conditions that initially buffer populations from collapse can simultaneously constrain their evolutionary response to emerging threats, and ultimately determine differential recovery following disease-induced declines.

## Main

Global environmental changes are reshaping selection pressures imposed on species^7–11^. Whether populations can adapt on a timescale necessary to avoid extinction is of critical importance given the pace of change^12^. Pathogen introductions can exert strong and often rapid selective pressure on wildlife populations^13^ and is thus a substantial challenge for species conservation in a rapidly changing world. However, population responses are often variable across the landscape following disease emergence, consisting of local extirpations, large-scale impacts, stabilization at lower densities, or rapid growth following declines^14–19^. Nevertheless, the conditions that lead to such divergent outcomes remain largely unknown. Understanding the circumstances that facilitate host resilience on ecologically relevant timescales is paramount to identify which populations are at the greatest risk of disease-driven declines and guide conservation action.

Historically, only ecological responses were considered rapid enough to prevent extinction in declining populations, but a growing body of evidence from field studies suggests that evolutionary responses can happen within a few generations^6,20–22^. If selective pressures are intense enough to act as a bottleneck and the appropriate amount of genetic variation and pre-adaptive traits are present in a population, then adaptation may occur fast enough to allow for population recovery^23–26^. This eco-evolutionary process, deemed “evolutionary rescue” (ER), has been proposed to occur in diverse systems in which populations undergo rapid environmental change^27–32^ (e.g. climate, invasive species, and pollution). Both pre-adaptive heterogeneity in hosts and abundance can affect species outcomes, such that larger more diverse populations are more likely to persist due to increased likelihood of phenotypes that will be robust to the emerging threats^21,33–34^. When hosts and pathogens impose reciprocal selection on one another within short timescales, these strong selective pressures can facilitate the evolution of potential adaptive strategies, such as pathogen resistance or infection tolerance^35^. Standing genetic variation is a crucial factor enabling host population evolution^3,23,36–37^, and each strategy results in different population-level processes with the potential to alter host-pathogen interactions over time^3,26,38,39^. Although infectious diseases can trigger sudden outbreaks and declines forcing host evolution^36^, it remains unclear why evolutionary rescue leads to coexistence in some local contexts and not others.

Gradients of selection, or ‘geographic mosaics’, may occur among populations following disease outbreaks due to variable host-pathogen interactions among locations^40^. If the intensity of selection varies environmentally, “hotspots”, where the reciprocal selection of host-pathogen coevolution is strong, and “coldspots”, where selection is weak, can occur^41^. The environment plays an important role in disease outcomes, such that coexistence may be modified by interactions with environmental conditions that lead to divergent host responses^3–5^. For example, variable environmental conditions across space and time have been shown to influence host susceptibility, disease severity, and transmission and infectivity which can scale to population-level effects^42–46^. Consequently, investigating the environmental influences that facilitate rapid evolution is critical to advance our understanding of differential host population outcomes following disease outbreaks.

White-nose syndrome (WNS), caused by the multi-host fungal pathogen *Pseudogymnoascus destructans*^47,48^, is a recently emerged disease of hibernating bats that has caused the collapse of North American bat populations. Both host and pathogen ecology drive infection patterns that lead to seasonal disease outbreaks during the hibernation period^49^. The temperature-dependent growth of *P. destructans* aligns with the conditions at which bats hibernate^50,51^, and thus, a drop in host body temperature coincides with reduced immune function that allows the pathogen to invade epidermal tissue and grow^47,52^. Following pathogen arrival, population responses have varied among host species and over space and time. Little brown bats (*Myotis lucifugus*), a long-lived previously common species, suffered severe initial declines (> 95%)^53,54^, yet several populations are now growing rapidly while others have just stabilized at low densities ^55,56^. Given that environmental conditions strongly influence pathogen replication rate, bats hibernating in warmer sites have been shown to experience more severe infections and impacts^43,57^. However, patterns of persistence have been less clear with variable recovery not aligning with initial expectations of species extinction^53^. The variability in impacts and recovery from WNS provides an opportunity to determine when and where evolutionary rescue is achieved and why.

Here we used two decades of population, environmental, and disease data to investigate drivers of recovery for *M. lucifugus* following WNS arrival. We paired this with a common garden experiment from a subset of these populations to explore differential selective pressures among remnant populations that have experienced variable degrees of declines and recovery. We translocated 120 *M. lucifugus* from eight hibernacula where WNS had been present for 8–16 years to common sites to control for environmental conditions, allowing us to disentangle the variability in resistance adaptation from initial selective pressures through survival and disease severity response. We hypothesized that variation in host adaptation results from a gradient of selection pressures that are largely driven by environmental conditions affecting the degree of initial decline severity.

## Results

### Decline severity and population recovery

We first explored factors influencing population dynamics of *M. lucifugus* over time since WNS arrival at 18 hibernacula across the Northeast and Midwest US. Initial declines were severe and occurred rapidly for almost all colonies (mean decline = 92% for 17 of 18 sites); however, some populations quickly stabilized (2–8 years following pathogen arrival) and began to recover to varying degrees (Fig. 1a, Extended Data Fig. 1). During the initial epidemic (1–4 years post arrival), environmental conditions imposed significant selective pressure on populations by influencing the severity of declines, such that warm sites experienced greater declines on average (Fig. 1b, Supplementary Table 1; p=0.057) and reached their minimum colony size prior to cold sites, which declined more slowly (p<0.001; Extended Data Fig. 2a, Supplementary Table 2). We found evidence that pre-WNS colony size played a role in the rate and severity of decline for cold sites, with larger populations in cold sites retaining more of their population throughout the initial epidemic, whereas warm sites experienced more severe declines regardless of colony size (Extended Data Fig. 2b, Supplementary Table 3). We categorized temperature across sites to reflect the dichotomy established with our common garden experiment, which used a smaller number of sites compared to population-level analyses, and results are presented as such for ease of interpretation. Continuous patterns generally align with categorical results, and all Supplemental Tables show results from both analyses.

**Figure 1.**
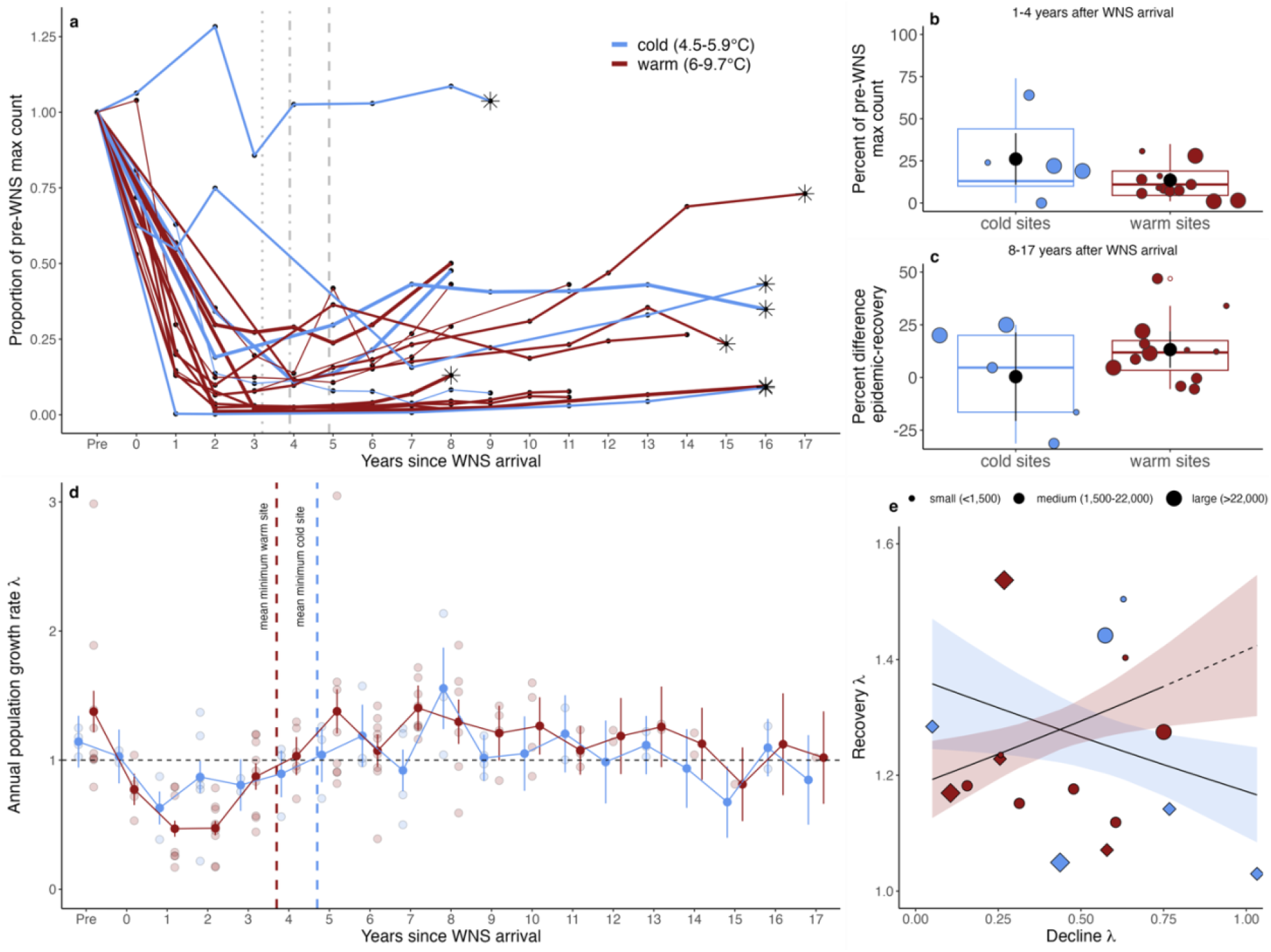
Recovery of little brown bats (*Myotis lucifugus*) following white-nose syndrome (WNS) arrival. (a) The proportion of the pre-WNS maximum (max) colony count over years since pathogen arrival (n=18 sites). Year = 0 represents the year in which the fungus was first detected. Line color represents the mean early winter hibernacula temperature (cold = <6°C, n =6 sites; warm = 6°C+, n=12 sites) and line thickness indicates the categorical pre-WNS population size (small: <1,500, n=4; medium: 1,500–22,000, n=9; large: 22,000–185,000, n=5; individual colony counts by year can be found in Extended Data Fig. 1). Vertical lines represent the mean year that colonies reached their minimum population size following pathogen arrival (dotted = large, 3.2 years; dot-dashed = medium, 3.9 years, dashed = small, 4.9 years). Stars indicate origin sites used in the common garden experiment. Variable population decline (b) and recovery (c) between cold and warm sites during the epidemic (1–4 years following pathogen arrival) and recovery (8+ years) periods. Declines are presented as the percent of the pre-WNS maximum count during the epidemic period, and recovery is shown as the percent difference between the epidemic and recovery periods. Black points show the model fitted mean percent and error bars represent the upper and lower confidence intervals. Boxplots show the inter-quartile range and whiskers correspond to smallest and largest values. (d) Annual population growth rates (*λ*) over years since pathogen arrival. Bold points and error bars represent model estimates ± standard error. The dashed line indicates a stable population growth rate (*λ*=1). Vertical lines represent the mean year warm (3.7 years), and cold (4.4 years) sites reached their minimum count. (e) Relationship between the decline (year of pathogen arrival to minimum colony count) and recovery (minimum colony count to most recent count) population growth rates for warm and cold sites. Diamonds indicate translocation sites and point size represents the categorical pre-WNS population size.

Recovery was also variable among colonies, ranging from just 2.4% to 73.1% of their pre-epidemic population size (Fig. 1a). In contrast to the pace and magnitude of initial declines, warm sites have grown faster since reaching their minimum and gained a larger percentage of their pre-WNS colony size during the recovery period (years 8–17, Fig. 1c). Colonies that were larger prior to WNS arrival experienced the greatest recovery (p<0.001, Supplementary Table 4). Annual population growth rates show that warm sites stabilized after reaching their minimum during the initial epidemic and are now consistently growing, suggesting that bats in these sites may be better adapted to surviving with WNS (*λ* > 1 after year 4 – except for year 15, only one data point), while cold sites experienced a shallower decline and less consistent subsequent growth rates (Fig. 1d, Supplemental Table 5). Overall, we find that the severity of initial declines interacted with environmental conditions to create a gradient of selection pressure that resulted in variable recovery across the continuum of colony sizes (Fig. 1e, Supplementary Table 6).

To test whether individuals in persisting colonies had variable resistance to WNS, we conducted a common garden experiment. We selected a subset of sites (eight) from the broader analysis (Fig. 1) that have persisting *M. lucifugus* colonies to serve as our origin sites. We selected these sites to encompass a gradient of hibernacula temperatures and pre-WNS population sizes (Extended Data Table 1, Extended Data Fig. 3 and Fig. 4, Supplementary Table 7). Additionally, these sites fall along a spectrum of initial decline severity and subsequent population growth. Given that pathogen growth is sensitive to environmental factors (temperature), we used two previously extirpated hibernacula, which are very warm and within in the range of optimal fungal growth, to serve as translocation sites as we predicted that the very warm temperatures in these sites would facilitate higher pathogen growth and thus allow us to compare resistance. We used fungal growth rate on bats and survival during the experiment as our response for measuring disease resistance.

Over the course of the experiment, on-host pathogen growth rates varied among and within origin sites, with individuals from site MW1 (no post-WNS decline) exhibiting the highest pathogen growth (Extended Data Fig. 5, Supplementary Table 8). We found that the severity of initial declines was correlated with the observed variability in host resistance, such that bats from populations with greater declines now experience significantly reduced pathogen growth (Fig 2a, p=0.022, Supplementary Table 9). Additionally, the variability we observed in host resistance was also driven by environmental conditions of origin sites; bats from warm sites had higher resistance (i.e. lower pathogen growth on average, p=0.015, Fig. 2b, Supplementary Table 9). Early winter fungal loads and body mass varied among origin sites at the onset of the experiment (Extended Data Fig. 7); however, over the hibernation season, 10% of individuals reduced their fungal load, including three bats from warm origin sites that completely cleared infection (Extended Data Fig. 5). Overall survival was 51% and varied by origin site (Figure 2c; range = 27–73%, Supplementary Table 10). Individuals from warm sites exhibited greater survival overall (p=0.021) with no effect of colony size. However, pre-WNS colony size significantly influenced outcomes for populations from cold sites, as we see the greater survival for bats originating in large cold sites compared to those from small cold sites (Fig. 2d, Supplemental Table 11). Survival was higher for bats at the Midwest translocation site, likely due to the difference in experimental time (i.e. 17 days shorter, p=0.052; Extended Data Fig. 6a, Supplemental Table 12), and for individuals from warm sites that entered the experiment with greater body mass and lower fungal loads (Extended Data Fig. 6b and Supplemental Table 13).

**Figure 2.**
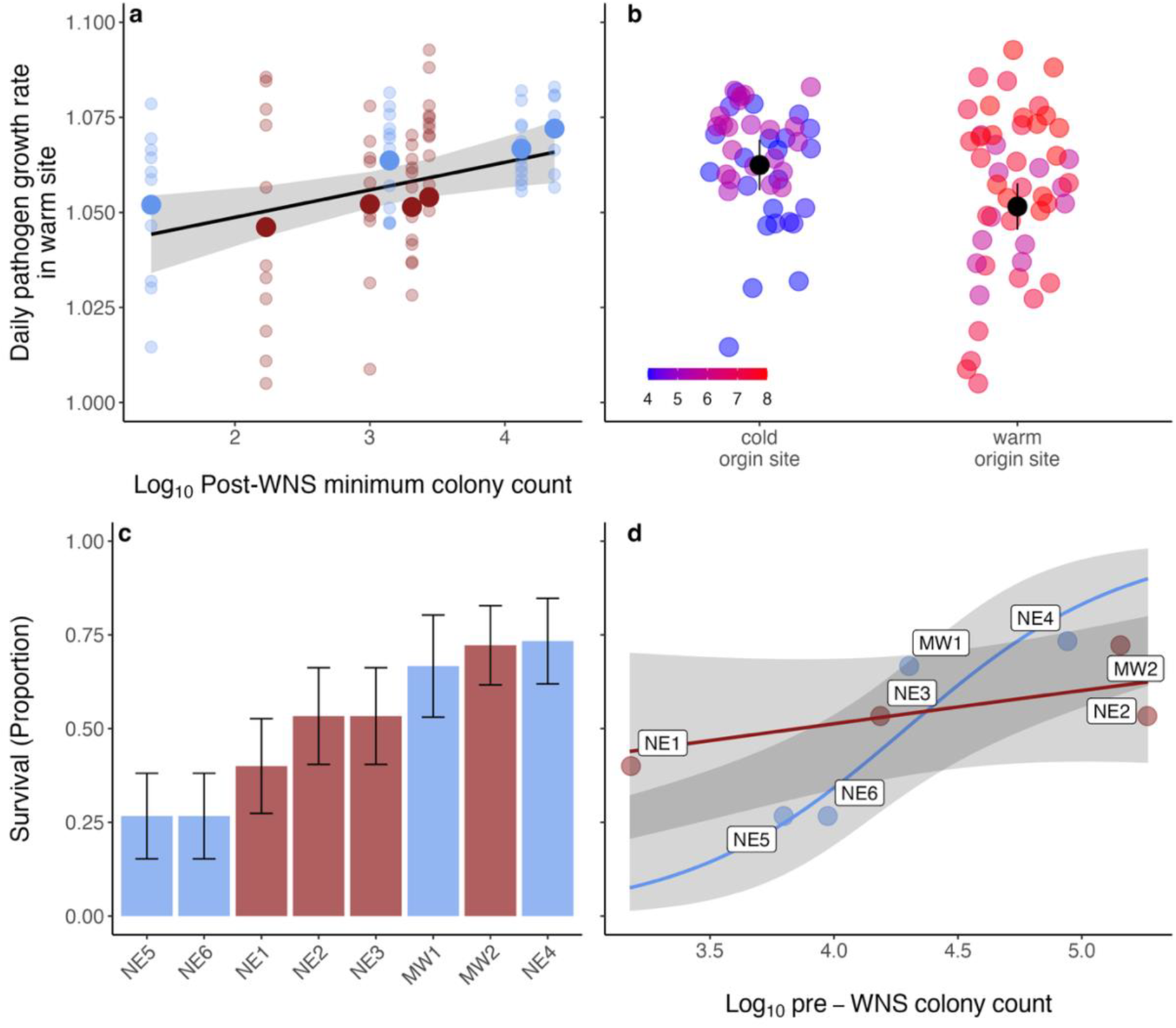
On-host pathogen growth and survival of translocated bats. The relationship between daily on-host pathogen growth rates of bats translocated to warm sites and (a) initial decline severity and (b) early winter temperature at origin sites. Point color depicts the early winter origin site temperature a) categorically (blue=cold, red=warm) and b) continuously (4.55-8.03°C). The solid black line indicates the model prediction, and the transparent ribbon represents the 95% confidence interval (a). Black points show the mean percent and error bars represent standard error (b). Three points have been removed (bats that cleared infection, daily growth rates of 0.97, 0.98, and 0.99) from both plots to improve visibility of the y-axis scale. (c) Proportion of bats from each origin site surviving the translocation experiment. Error bars represent standard error. (d) Relationship between survival, the pre-WNS colony count, and early winter temperature at origin sites. Colored lines indicate the model predictions for cold vs warm sites, and the transparent ribbon shows the 95% confidence intervals. Bars (c) and points (d) are colored according to their early winter origin site temperature.

Finally, to assess changes in resistance of natural populations over time, we used field data collected from historic populations during the initial epidemic (years 1–4 following pathogen arrival; n=872 individuals from 25 sites across the Northeast and Midwest) and from contemporary populations (8–16 years following pathogen arrival) sampled at the conclusion of our experiment (non-experimental free-flying bats within 7 origin sites, n = 105 individuals) to compared infection intensity (measured as late hibernation fungal loads). We found that historic and contemporary populations in cold sites show little change over time and comparable fungal loads (Fig. 3a,c, & d; p=0.939), whereas fungal loads for contemporary populations in warm sites are an order of magnitude lower compared to the epidemic period (Fig. 3b,c, & d; p < 0.001; Supplementary Table 14). Additionally, we compared infection intensity between free-flying bats sampled within origin sites and experimental bats to explore how fungal loads varied when individuals from contemporary populations were challenged. Bats originating from cold sites had the highest fungal loads when translocated to warm sites, while free-flying bats from warm sites had the lowest fungal loads (Fig. 3e & f; Supplementary Table 15). Translocation sites were 2.6°C (±0.69 sd) and 0.5°C (±0.65 sd) warmer on average than cold and warm origin sites, respectively (Fig. 3g). Free-flying bats from cold sites had significantly higher degrees of tissue invasion, indicative of severe disease, compared to free-flying bats from warm sites (Extended Data Fig. 8, Supplementary Table 16).

**Figure 3.**
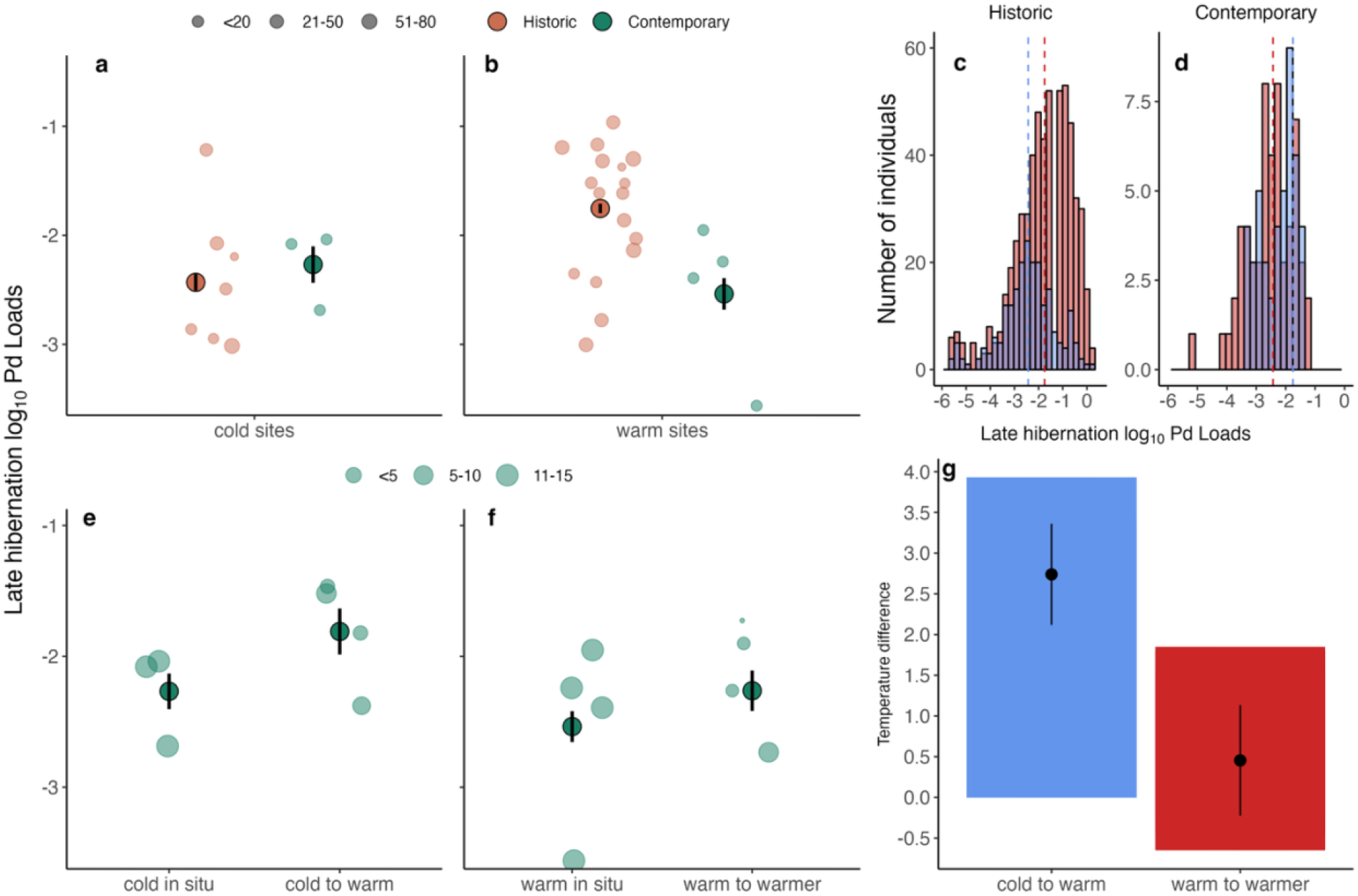
Mean infection intensity over time since pathogen arrival. (a–d) Historic (years 1–4) and contemporary (years 8–16) late hibernation fungal loads (*Pseudogymnoascus destructans*, Pd) of *Myotis lucifugus* in cold (n=10) and warm (n=22) hibernacula (n=977, mean=35, range=1–80 bats sampled per site). Size of colored points represents the number of bats sampled per site. Bold points and error bars represent model estimates ± standard error. (e–f) Contemporary late hibernation fungal loads for free-flying bats in cold (cold *in situ*, n=3) and warm (warm *in situ*, n=4) hibernation sites (n=105 individuals), and bats in warm translocation sites (cold to warm, warm to warmer; n=4 sites of each temperature category; n=62 individuals). Size of colored points represents the number of bats sampled per site. Bold points and error bars represent model estimates ± standard error. (g) Temperature difference between cold and warm origin sites and the temperature of the translocation sites. Translocation sites averaged 2.6°C warmer than cold origin sites and 0.5°C degrees warmer than warm origin sites.

### Rapid adaptation drives population recovery

While all species experience selective pressure from interactions with their biotic and abiotic environments^58^, the pressure exerted by emerging pathogens can be strong enough to induce an evolutionary response that facilitates host adaptation^,39^. However, the outcomes of host-pathogen interactions can vary from host recovery to extinction, and thus understanding what drives variable population responses is vital for promoting species persistence. Our results show that the environmental conditions that helped prevent population collapse simultaneously hindered recovery and adaptation. Additionally, we illustrate that rapid host adaptation is possible for a long-lived species with long generation times following novel pathogen emergence.

Rapid and precipitous declines at most sites during the WNS epidemic represent a disease-induced bottleneck and gradient of selective pressures that are ultimately driven by variable environmental conditions (Fig. 4). Warm hibernacula appear to be hotspots of adaptation where the intensity of selection from the pathogen on hosts is greatest. In contrast, populations originating from cold hibernacula (coldspots) are initially buffered from severe declines and thus experience weak selective pressure due to environmentally-mediated reductions in pathogen growth, and consequently, these conditions have hindered recovery. Contemporary populations originating in warm sites have significantly reduced infection intensity in late hibernation compared to the initial epidemic period, and the ability of individuals to reduce and clear infections suggests the rapid evolution of resistance traits. These processes appear to have led to local host adaptation and divergent population responses just a decade after disease emergence.

**Figure 4.**
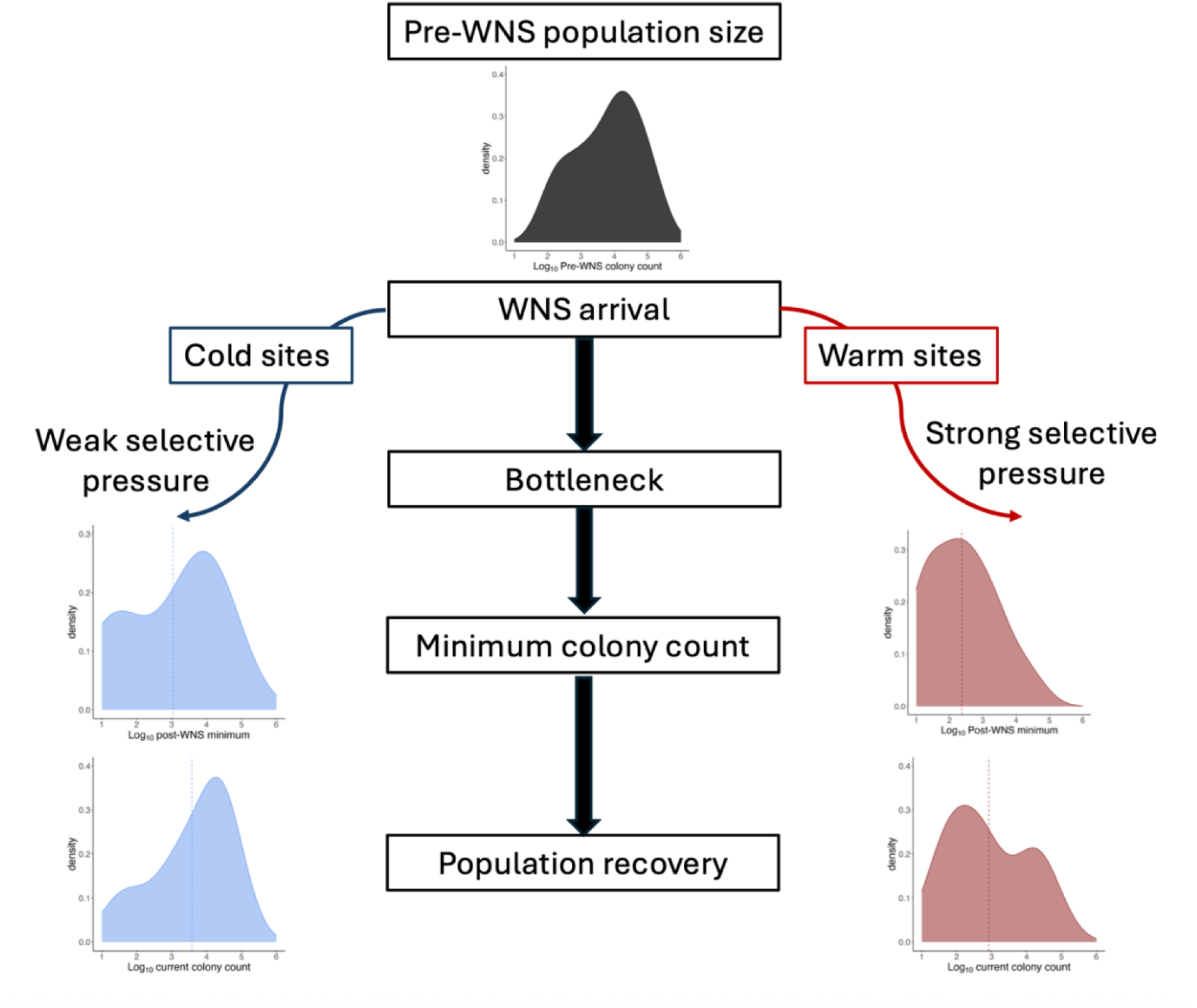
Theorized pathway of differential population recovery and adaptation following pathogen introduction. Following the arrival of white-nose syndrome (WNS), environmental conditions, specifically early hibernation temperatures, created a gradient of selection on the 18 sites we examined, in which populations in cold (n=6) versus warm sites (n=12) experienced differential selective pressures resulting from environmentally mediated pathogen growth. Rapid and precipitous declines imposed a bottleneck, with declining populations reaching their minimum size within 4 years, on average. Populations in warm sites experienced strong selective pressure to adapt likely due to greater pathogen growth. Populations in cold sites were initially buffered from severe declines due to reduced pathogen growth, and although they retained a greater proportion of their pre-WNS population, these conditions have now hindered recovery. Density plots show the colony counts for sites at their post-WNS minimum size and their most recent counts. Dotted vertical lines indicate the mean colony count.

We find that environmental conditions interacted with pre-epidemic colony size to influence the rate and severity of declines as well as subsequent recovery and survival. Populations rapidly reached their minimum size following pathogen arrival; however, declines were more gradual for small and/or cold sites. For example, populations in small cold sites took 7 years on average to reach their minimum size, while populations in large warm size reached their minimum in 3 years on average. Less than a decade after pathogen arrival, we see consistently positive growth rates for populations in warm sites, whereas those occurring in cold sites are lower on average. Additionally, populations that were large prior to pathogen arrival now exhibit greater increases in colony size and/or growth rates during the recovery period, while small populations remain at low abundance. Our results are consistent with the theory of evolutionary rescue (ER), indicating that initial population size is a primary determinant of the ability of a population to adapt successfully to environmental change^21,33,34^. Large populations are more likely to contain enough individuals robust to new environmental conditions and make it through population bottlenecks, while small colonies may lack the necessary diversity or pre-adaptive traits to respond rapidly^59^, leading to stagnate recovery. The more time spent below a critical threshold of abundance reduces the degree of ER and increases vulnerability to extirpation due to stochasticity or Allee effects^24,26,60,61^. In our results we see the greatest recovery for populations that experience the strongest initial selective pressure (i.e., warm sites that reached their minimum colony size early on during the epidemic). Although bats originating from warm sites exhibit the greatest survival in our common garden experiment, pre-epidemic population size plays a larger role to promote survival of bats from cold sites. Large populations, regardless of environmental conditions, may contain individuals with resistant alleles that are more strongly selected in warm sites yet remain in cold sites through the bottleneck, aligning with ER theory. While empirical support for ER remains rare for wild vertebrate populations as most cases are restricted to short-lived fecund species with large population sizes^62^, our study adds a significant contribution to the growing body of evidence showing rapid adaptation can occur for hosts across diverse disease systems, such as amphibians and chytridiomycosis^63^, chronic wasting disease in cervids^64^, and prairie dogs affected by plague epidemics^65^.

### Environmental conditions shape selective pressures to drive variability in host resistance

The environmental dependence of pathogen growth can affect the progression of an epidemic in complex ways, by creating hotspots or cold spots of pathogen transmission^66^, host susceptibility^67^, or disease severity^68^, particularly for multi-host pathogens with variable host-species responses under diverging environmental conditions^69^. Temperate bats select winter roost sites with conditions that maximize energy savings and reduce evaporative water loss^50^; however, warmer hibernacula temperatures are highly conducive for growth of *P. destructans*^51^, and because the roosting microclimates that bats select largely reflect the thermal conditions necessary or favorable for pathogen growth, environmental conditions can have a large effect on individual infection severity and survival^57^. Prior to WNS arrival, colder sites likely provided high-quality habitat that supported greater metabolic efficiency during hibernation. Previous studies have found that these conditions provide thermal refugia that reduced infection severity and survival following WNS arrival for *M. lucifugus*, while warm sites initially acted as ecological traps facilitating high pathogen growth and disease severity^57^, and thus increased decline severity^54^. While our data supports this pattern (i.e., severe infections and population declines at warm sites during the initial epidemic), we show that these same conditions ultimately impose a gradient of selective pressures on populations that lead to opposing forces during the decline and recovery phases.

Here we present several lines of evidence supporting evolution of resistance mechanisms within populations of *M. lucifugus*. Pathogen growth rates differed based on early hibernation temperatures, such that bats originating from warm sites experienced lower on-host growth rates in our common garden. Additionally, populations that suffered the most severe disease-induced declines displayed reduced infection severity, and the ability of individuals to reduce or clear infections over the course of hibernation suggests the evolution of traits conferring resistance. Interestingly, we observed greater variability in on-host pathogen growth rates for populations from warm sites, and individuals that reduced or cleared infections did not necessarily originate from sites with the highest survival. This suggests there may be a tradeoff occurring where the adaptive mechanism is costly, as not all individuals within specific populations are resistant. This pattern aligns with previous studies that demonstrate intensity-reduction resistance strategies lead to faster recovery for host populations^26^, yet resistant alleles are unlikely to reach fixation due to trade-offs that promote polymorphisms in resistant traits and subsequently relax positive selection^35^. Historic populations exhibited similar infection severity regardless of environmental conditions, yet we observed a markedly different outcome for contemporary populations, based on early winter hibernacula temperature—bats that originated in warm sites had significantly reduced infection severity compared to the initial epidemic period, even when challenged in warmer sites. Host susceptibility and disease severity were higher for individuals occurring in cold spots when challenged with warm conditions, likely due to weak selective pressure in cold hibernacula during the epidemic. Our results support prior studies that identified genomic signatures of selection for *M. lucifugus*^70,71^ and highlight the importance of standing genetic variation for rapid host responses^23^. Nevertheless, further research is warranted to provide a deeper understanding of the tradeoffs of specific strategies that support persistence and lead to differential host outcomes.

### Conservation in a rapidly changing world

Successful conservation of host populations will require flexible strategies that reflect the complex nature of emerging infectious diseases. Although our study strongly suggests that resistance offers protection from severe disease outcomes in some contemporary populations of *M. lucifugus*, recovery has been variable; thus, our results have important implications for management of WNS-impacted bats. Environmental treatments and habitat modifications have been explored as management tools to increase survival by reducing the environmental pathogen reservoir and creating conditions that reduce fungal growth^72,73^. However, treatment efforts may be counterproductive if they interfere with host’s adaptive responses for populations already in recovery or lead to maladaptations under conditions that impose weak selective pressure^64,74^. Small colonies remain vulnerable, particularly those occurring in environments that experience weak selective pressure (i.e. cold sites) or in areas where declines are very high leading to stochastic loss of resistance alleles, as predicted by evolutionary rescue theory, which potentially aligns with reports of slower recovery in bat populations in the southern U.S.^56^.

Significant environmental changes, such as the emergence of new pathogens, can cause widespread population losses and result in host extinction, but may also impose strong selective pressure that facilitates recovery. As emerging diseases continue to pose a substantial threat to global health and biodiversity, understanding how populations respond is imperative for effective management. Documenting disease-driven selection pressure remains challenging due to the complexity of host-pathogen-environment interactions that shape population responses. Host outcomes will depend on the structure of selective pressures acting on host-pathogen-environment interactions, and we show that local host adaptation leads to individual outcomes that scale up to population-level effects to drive variable recovery among populations.

## Methods

### Data collection

#### Population data

We analyzed population count data for 18 colonies of hibernating *M. lucifugus* from four US states, collected between 1984 and 2024. Surveys were conducted via visual counts by state and federal agencies or the authors. For each colony, the year of detection of *P. destructans* infection within the hibernaculum was recorded and used to assign the year of pathogen arrival (YOA). We used the most recent census to represent colony size before the onset of WNS infection (pre-WNS maximum colony count, mean=4.8 years before pathogen arrival, range: 1–25 years). Environmental data (site temperatures) were collected at each hibernaculum with various data loggers between 2001–2024. Hibernacula were classified as cold (4.5–5.9ºC, n=6 sites) or warm sites (6–9.7ºC, n=12 sites) according to the mean early winter temperature (November–December). We categorized the temperature variable to align with the formation of our common garden experiment, which used a smaller number of sites compared to population-level analyses, and we used a cutoff of 6ºC between cold and warm sites to reflect the distribution of winter temperatures at origin sites. For statistical analyses incorporating the categorical early winter hibernation temperature as a predictor variable, we also ran models with the same structure using the continuous early winter hibernation temperature to show the relationship directionality for both predictor types (results for both categorical and continuous temperature models are found in Supplemental Tables 1–16). The mean early winter temperature was selected as an environmental variable of importance because pathogen prevalence and infection intensity typically spike shortly after bats enter hibernation, and thus, this measure appears the be the most influential environmental metric affecting disease progression^69^. Colonies were classified into population size categories according to the interquartile of their pre-WNS maximum colony count (small = <1,500, medium = 1,500–22,000, large = >22,000– 185,000). We estimated the mean year that colonies of each population-size category and temperature category reached their minimum count following pathogen arrival. Lastly, we classified colony count data by epidemic phase, with the ‘pre-invasion’ phase defined as epidemic years ≤0, ‘epidemic’ years one through four, and ‘recovery’ defined as epidemic years ≤5. Phases were established based on the mean year that all populations within our dataset reached their minimum count (3.9 years), as well as previous results demonstrating populations approaching stability by year 4^53,75,76^.

### Translocation ‘Common Garden’ experiment

We translocated 120 male bats from eight hibernacula of persisting little brown bat colonies (origin sites) that have experienced differential effects of disease within two regions of the US, the Northeast (NE) and the Midwest (MW), to two hibernacula previously extirpated by white-nose syndrome in Vermont (NE) and Wisconsin (MW). Selected origin sites fall along a spectrum of initial population declines and subsequent population growth, encompassing a range of hibernacula temperatures and pre-WNS population sizes (Extended Data Table 1, Extended Data Fig. 3 and 4). Origin sites were classified as cold (< 6°C, n=4) or warm (≥ 6°C, n=4) according to early winter hibernation temperatures (range=4.55–8.03°C). Ninety bats were transported from six hibernacula in New York (NE1–6) to one hibernaculum in Vermont (NE-T), and 30 bats were moved from two hibernacula, one in Michigan (MW1) and one in Wisconsin (MW2), to one hibernaculum in Wisconsin (MW-T; Extended Data Table 1). We selected two “garden” translocation sites to limit disturbance and transportation time for bats. This design allowed us to incorporate a population of bats that has yet to experience declines from WNS (MW1), despite detection of *P. destructans* (eight years since pathogen arrival), without causing excessive stress to most study animals. Prior to hibernation, we installed 18 (NE) and six (MW), four-sided reptile cages (dimensions: 12”W, 18”L, 20”H) in historic roosting locations within each extirpated site. We mounted each cage so that the open side was in contact with the wall or ceiling to allow bats to roost on the substrate. Contact points between the cage and the substrate were sealed to prevent bat escape and eliminate the possibility of predation. Water was provided in poultry waterers for each cage.

In early hibernation (NE: October 25, 2023; MW: November 1, 2023), we collected 15 bats from each NE origin site (NE1–6), 12 bats from MW1, and 18 bats from MW2. Sample sizes were determined by a power analysis and represent the minimum number of bats per origin site necessary to detect a significant effect (p < 0.05) of selection in relation to population survival estimates and post-WNS population growth rates at each origin site. Each bat received a unique forearm band (2.9 mm; Porzana, Icklesham, UK), and individuals were randomly assigned to a cage within their respective translocation sites. Bats were randomly mixed among cages at their respective translocation sites to ensure all individuals were adequately infected at the onset of the experiment (five individual bats per cage). We placed temperature loggers (Maxim Integrade iButtons) within each cage to record roosting temperatures, and we installed psychrometers within the general cage area of each translocation hibernacula to record temperature and relative humidity (Extended Data Table 1 and Figure 3). Mean cage roosting temperatures were 7.56°C (range=7–8.5°C) and 8.69°C (range=5.8–10°C) at NE-T and MW-T, respectively. Both translocation sites exhibited 100% average relative humidity throughout the hibernation season, with vapor pressure deficit ranging between −0.021–0.024 (NE-T) and −0.091–0.008 (MW-T).

We placed each bat in a cloth bag and transported the animal in a temperature-controlled cooler (kept at a mean of 7°C) to translocation sites so that the bats would remain in torpor and limit energy expenditure^38^. We replicated the transportation disturbance across all treatment groups by keeping bats from each origin site in the coolers for the same amount of time. In mid-hibernation (NE translocation site: 57 days; MW translocation site: 66 days), each translocation site was visited to record survival status of the bats through visual inspection and collect epidermal swab samples from each bat and the cage substrate. In March 2024, 149 (NE) and 132 (MW) days following initial translocation, we returned to each translocation site to collect data and terminate the experiment. We released all surviving individuals from NE1–6 and MW2 at their site of origin; all bats from MW1 and those from MW2 that had been caged with MW1 individuals were humanely euthanized. For all survival analyses, we tested translocation site as an additive effect to determine if the difference in experimental time influenced survival; however, all models indicated no significant differences due to translocation site, and thus, this variable was excluded from final models.

We measured three disease severity metrics—pathogen growth rate, UV wing fluorescence, and body condition—for each bat at various times throughout the experiment. Epidermal swab samples were collected during early hibernation (onset of the experiment), mid-hibernation, and late hibernation (termination of the experiment) with a standardized swabbing technique^48^. To measure the on-host pathogen growth rate of *P. destructans* across the hibernation season, we used real-time polymerase chain reaction (qPCR) to quantify fungal loads at each time point^77^. We weighed each bat at the onset and termination of the experiment using an electronic scale, and calculated weight loss over the course of hibernation by subtracting the final hibernation weight of each individual from hibernation weight at the time of collection. At the termination of the experiment, we collected data on the severity of tissue invasion by transilluminating both wings of each individual using a 9-watt 368-nm ultraviolet light and calculating the average proportion of both wings displaying orange fluorescence^38^. At the termination of the experiment, we collected epidermal swab samples, weights, and UV wing photos from 15 non-experimental free-flying bats at 7 of the origin sites, to compare with caged individuals and represent late hibernation infection severity of contemporary populations. Additionally, swab samples from late hibernation were collected from bats (n=872 samples, mean bats sampled per site = 35, range = 1–80) at 25 hibernacula across 4 states in the Northeast and Midwest US between 2011 and 2021 (1–4 years following pathogen arrival) to represent infection severity of historic populations^69,76^. All bat handling and housing procedures followed the approved IACUC protocol (#23-206) through Virginia Tech, and decontamination followed US Fish and Wildlife Service protocols, as well as state recommendations.

### Statistical analyses

#### Population declines and recovery

We conducted all analyses using package glmmTMB^78^ in R v4.3.1. Model descriptions (Models 1–16) are shown in Supplemental Table 17. For any model incorporating the categorical early winter hibernation temperature as a predictor variable, we also ran an additional model with the same structure using the continuous early winter hibernation temperature to show the relationship directionality for both predictor types. To explore the change in colony counts over time, we calculated the proportion of the pre-WNS colony count that remained in each year since pathogen arrival (YSW). Origin site MW1 was removed from the dataset for Models 1–4, as it has not experienced any population decline post-WNS arrival^79^. We characterized population recovery by calculating the percentage of the pre-WNS colony count that remained in the epidemic period (years 1–4 following pathogen arrival), and the difference between the percent remaining during the epidemic and the recovery period (years 8–17). To explore associations between environmental conditions within hibernacula, pre-WNS population size, and the response of colonies to WNS, we used a series of generalized linear models. First, to test for the effect of early winter hibernation temperature on initial declines, we used a gamma distribution with log link function and the percent of the pre-WNS colony size remaining during the epidemic as the response variable (Model 1). We then tested whether the early winter hibernation temperature, log_10_ pre-WNS colony size, and their interaction influenced 1) the years to reach the post-WNS minimum count (Model 2 - Poisson distribution with log link function), 2) the percent of the pre-WNS population remaining at the minimum (Model 3 - gamma distribution with log link function), and 3) the percent difference of the pre-WNS population remaining during the epidemic and recovery years (Model 4 - negative binomial distribution with log link function).

We estimated annual population growth rates, *λ*, using the pre-WNS maximum colony size and all subsequent colony counts (mean = 6.8 surveys per site post-WNS arrival, ranging 8– 17 YSW). Because surveys were not conducted every year after WNS detection, we calculated the average yearly population growth rate, *λ*, following the arrival of WNS by adjusting for the number of years between WNS detection at a hibernaculum and the post-WNS colony count^53^. We then modeled the interaction between early winter hibernation temperature and years since WNS arrival (YSW; categorical) on annual population growth rates using a gamma distribution with log link function (Model 5). We explored the relationship between decline severity and subsequent population growth by calculating the population growth rate (*λ*) during the decline (year of WNS arrival to the year of the WNS minimum colony count) and the recovery period (year of the WNS minimum to most recent colony count) for each site following the same method described above to account for gaps in survey years. We then modeled the effect of the decline growth rate on the recovery growth rate using a generalized linear model with a gamma distribution (log link function) and early winter hibernation temperature as a fixed effect (Model 6), and then ran the same model without temperature for just the translocation site populations (Model 7).

### Disease severity and survival

For bats used in the translocation experiment, we calculated daily on-host pathogen growth rates across the hibernation period using our estimates of fungal loads during early hibernation (N_*0*_), fungal loads during late hibernation (N_*t*_), and the number of weeks between collection of the early and late samples (*t*)^69^. We compared daily pathogen growth rates for bats based on origin site with a generalized linear mixed model (gamma distribution, log link) and cage ID as a random effect (Model 8). To examine the effect of early winter hibernation origin site temperature and the post-WNS minimum colony count on daily pathogen growth rates, we used a generalized linear mixed model (gamma distribution, log link function) with cage ID as a random effect (Model 9).

We fit a generalized linear mixed model (binomial distribution, logit link) with survival as the response variable, origin site as a fixed effect, and cage ID as a random effect to assess how survival varied by origin site (Model 10). To explore the effects of decline severity and pre-WNS population size on survival, we fit a binomial (logit link) generalized linear mixed models with cage ID as a random effect and the pre-WNS colony count, early winter temperature, and their interaction as predictors (Model 11). We used a generalized linear mixed model (binomial distribution, logit link) to test the difference in survival between translocation sites, with cage ID as a random effect (Model 12). Because body condition and infection status at the onset of hibernation may influence WNS-induced mortality, we examined the effect of early hibernation fungal load, early hibernation body mass, and early hibernation origin site temperature on survival using a binomial generalized linear mixed model (logit link) with cage ID as a random effect for all individuals that entered the experiment *P. destructans* positive (Model 13).

### Infection intensity patterns over time

We compared infection intensity (measured as log_10_ late hibernation fungal loads) between non-experimental free-flying bats in contemporary populations sampled at the conclusion of our experiment (sampled within 7 origin sites) and historic populations sampled during the initial epidemic, with a generalized linear model and early winter hibernation temperature, phase (historic versus contemporary), and their interaction as fixed effects (Model 14). To explore the difference between infection intensity of contemporary caged versus free-flying bats, we used a generalized linear model with status (caged versus free-flying), early winter hibernation temperature, and their interaction as fixed effects (Model 15). Lastly, we compared tissue invasion (measured as the proportion of infected wing tissue, indicated by orange fluorescence) between caged and free-flying bats with a generalized linear model (gamma distribution, log link) and early winter hibernation temperature as a fixed effect (Model 16).

## Data and code availability

The datasets generated in this study and all code needed to reproduce the analyses and figures of this manuscript are available at Github (https://github.com/s02hoff/MYLUrecovery). Site names are anonymized to protect endangered species and landowners.

## Acknowledgements

We thank the University of Wisconsin Milwaukee Field Station, the U.S. Forest Service, and Consumers Energy for site access.

## Funding

Funding for this work was provided by NSF grant no. DEB-1911853 to KEL and JRH.

## Author contributions

This study was conceptualized by K.E.L., J.R.H., and S.H. All authors contributed to the data collection (S.H., J.R.H., J.P.W., H.M.K., J.A.R., J.E.D., A.B.B., A.M., A.K., A.T.G., M.J.K., N.A.L., C.D.K., K.E.L.). Data manipulation and analyses were performed by S.H. (with conceptual and technical input from K.E.L. and J.R.H.). S.H. wrote the manuscript, K.E.L. and J.R.H. edited the manuscript. All authors reviewed the final version.

## Notes

### Competing Interest Statement

The authors have declared no competing interest.

