## Supplemental Tables for "Environmental conditions drive selection and recovery following disease-induced declines"

Supplementary Table 1.

| **Figure 1b, Model 1 - Response variable**: percent of pre-WNS population size remaining during the epidemic (years 1-4) | | | |
| --- | --- | --- | --- |
| **Generalized linear model with gamma distribution and log link function:** Percent ~ (early winter temperature) | | | |
| N=17 populations (site MW1 removed, no decline) | | | |
| *Term* | *Estimate* | *Std. error* | *P value* |
| Categorical temperature model – reference level: cold | | |  |
| Intercept | 3.280 | 0.272 | <0.001 |
| tempcat:warm | -0.629 | 0.317 | 0.054 |
| Continuous temperature model | | |  |
| Intercept | 3.658 | 0.655 | <0.001 |
| origin_temp | -0.114 | 0.089 | 0.199 |

Supplementary Table 2.

| **ED Figure 2a, Model 2 - Response variable**: years to WNS minimum colony count  **Poisson generalized linear model with log link function:** Years to WNS min ~ (early winter hibernation temperature) * Log_10_(Pre-WNS colony count) | | | |
| --- | --- | --- | --- |
| N=17 populations (site MW1 removed, no decline) | | |  |
| *Term* | *Estimate* | *Std. error* | *P value* |
| Categorical temperature model – reference level: cold | | | |
| Intercept | 3.142 | 0.327 | <0.001 |
| tempcat-warm | -1.465 | 0.376 | <0.001 |
| Precountlog | -0.422 | 0.087 | <0.001 |
| tempcat-warm:Precountlog | 0.320 | 0.101 | 0.002 |
| Continuous temperature model | | | |
| Intercept | 4.206 | 0.885 | <0.001 |
| origin_temp | -0.311 | 0.121 | 0.010 |
| Precountlog | -0.762 | 0.234 | 0.001 |
| origin_temp:Precountlog | 0.084 | 0.032 | 0.009 |

Supplementary Table 3.

| **ED Figure 2b, Model 3 - response variable**: proportion of pre-WNS count at the minimum | | | |
| --- | --- | --- | --- |
| **Generalized linear model with gamma distribution and log link function:** (prop at min) ~ (early winter temperature) * Log_10_ (pre-WNS max population size) | | | |
| N=17 populations (site MW1 removed, no decline) | |  |  |
| *Term* | *Estimate* | *Std. error* | *P value* |
| Categorical temperature model – reference level: cold | | | |
| Intercept | -5.087 | 0.667 | <0.001 |
| Precountlog | 0.663 | 0.166 | <0.001 |
| tempcat:warm | 3.008 | 0.739 | <0.001 |
| precountlog-tempcat:warm | -0.812 | 0.186 | <0.001 |
| Continuous temperature model | | | |
| Intercept | -3.244 | 1.518 | 0.034 |
| Precountlog | 0.268 | 0.386 | 0.489 |
| origin_temp | 0.085 | 0.206 | 0.680 |
| Precountlog:origin_temp | -0.034 | 0.053 | 0.524 |

Supplementary Table 4.

| **Figure 1c, Model 4 - response variable**: percent difference of the pre-WNS population size remaining between (years 1-4) and (years 8-17) | | | |
| --- | --- | --- | --- |
| **Generalized linear model with gamma distribution and log link function:** (Percent difference) ~ (early winter hibernation temperature) * Log_10_(Pre-WNS colony count) | | | |
| N=17 populations (site MW1 removed, no decline) | | |  |
| *Term* | *Estimate* | *Std. error* | *P value* |
| Categorical temperature model – reference level: cold | |  |  |
| Intercept | -1.168 | 0.788 | 0.138 |
| Precountlog | 0.956 | 0.197 | <0.001 |
| tempcat-warm | 4.384 | 0.850 | <0.001 |
| Precountlog:tempcat-warm | -1.019 | 0.214 | <0.001 |
| Continuous temperature model | |  |  |
| Intercept | -0.920 | 2.033 | 0.651 |
| origin_temp | 0.496 | 0.268 | 0.065 |
| Precountlog | 0.968 | 0.491 | 0.049 |
| origin_temp:Precountlog | -0.125 | 0.065 | 0.055 |

Supplementary Table 5.

| **Figure 1d, Model 5 - response variable**: annual population growth rate (𝜆) | | |  |
| --- | --- | --- | --- |
| **Generalized linear model with gamma distribution and log link function:** (lambda) ~ (early winter hibernacula temperature) * (year since WNS arrival[categorical]) | | | |
| N=18 populations |  |  |  |
| *Term* | *Estimate* | *Std. error* | *P value* |
| Categorical temperature model – reference level: cold | |  |  |
| Intercept | 0.134 | 0.175 | 0.447 |
| tempcat-warm | 0.185 | 0.211 | 0.381 |
| YSW:0 | -0.106 | 0.268 | 0.694 |
| YSW:1 | -0.597 | 0.268 | 0.028 |
| YSW:2 | -0.273 | 0.227 | 0.230 |
| YSW:3 | -0.348 | 0.304 | 0.254 |
| YSW:4 | -0.246 | 0.268 | 0.361 |
| YSW:5 | -0.094 | 0.268 | 0.727 |
| YSW:6 | 0.039 | 0.268 | 0.884 |
| YSW:7 | -0.216 | 0.248 | 0.385 |
| YSW:8 | 0.309 | 0.268 | 0.252 |
| YSW:9 | -0.117 | 0.248 | 0.639 |
| YSW:10 | -0.084 | 0.210 | 0.691 |
| YSW:11 | 0.052 | 0.304 | 0.864 |
| YSW:12 | -0.148 | 0.274 | 0.591 |
| YSW:13 | -0.023 | 0.268 | 0.931 |
| YSW:14 | -0.199 | 0.274 | 0.468 |
| YSW:15 | -0.526 | 0.370 | 0.158 |
| YSW:16 | -0.041 | 0.268 | 0.879 |
| YSW:17 | -0.299 | 0.370 | 0.420 |
| tempcat-warm:YSW0 | -0.470 | 0.332 | 0.159 |
| tempcat-warm:YSW1 | -0.480 | 0.321 | 0.138 |
| tempcat-warm:YSW2 | -0.794 | 0.280 | 0.005 |
| tempcat-warm:YSW3 | -0.107 | 0.346 | 0.758 |
| tempcat-warm:YSW4 | -0.042 | 0.326 | 0.897 |
| tempcat-warm:YSW5 | 0.095 | 0.318 | 0.764 |
| tempcat-warm:YSW6 | -0.288 | 0.315 | 0.363 |
| tempcat-warm:YSW7 | 0.237 | 0.301 | 0.433 |
| tempcat-warm:YSW8 | -0.367 | 0.321 | 0.255 |
| tempcat-warm:YSW9 | -0.013 | 0.326 | 0.969 |
| tempcat-warm:YSW10 | NA | NA | NA |
| tempcat-warm:YSW11 | -0.297 | 0.370 | 0.423 |
| tempcat-warm:YSW12 | NA | NA | NA |
| tempcat-warm:YSW13 | -0.066 | 0.384 | 0.863 |
| tempcat-warm:YSW14 | NA | NA | NA |
| tempcat-warm:YSW15 | NA | NA | NA |
| tempcat-warm:YSW16 | -0.162 | 0.457 | 0.724 |
| tempcat-warm:YSW17 | NA | NA | NA |
| Continuous temperature model | |  |  |
| Intercept | -0.539 | 0.445 | 0.228 |
| origin_temp | 0.112 | 0.062 | 0.073 |
| YSW:0 | 1.028 | 0.686 | 0.137 |
| YSW:1 | 0.738 | 0.642 | 0.253 |
| YSW:2 | 1.177 | 0.604 | 0.054 |
| YSW:3 | 0.960 | 0.781 | 0.222 |
| YSW:4 | 0.108 | 0.742 | 0.885 |
| YSW:5 | 0.094 | 0.704 | 0.895 |
| YSW:6 | 1.096 | 0.665 | 0.102 |
| YSW:7 | -0.080 | 0.626 | 0.899 |
| YSW:8 | 1.585 | 0.683 | 0.022 |
| YSW:9 | 0.415 | 0.669 | 0.537 |
| YSW:10 | -2.018 | 2.112 | 0.341 |
| YSW:11 | 1.045 | 0.828 | 0.210 |
| YSW:12 | 0.097 | 4.052 | 0.981 |
| YSW:13 | 0.546 | 0.935 | 0.561 |
| YSW:14 | -0.580 | 4.052 | 0.886 |
| YSW:15 | -0.505 | 0.361 | 0.165 |
| YSW:16 | 1.013 | 1.415 | 0.476 |
| YSW:17 | -0.307 | 0.363 | 0.398 |
| origin_temp:YSW0 | -0.201 | 0.094 | 0.034 |
| origin_temp:YSW1 | -0.236 | 0.089 | 0.009 |
| origin_temp:YSW2 | -0.282 | 0.086 | 0.001 |
| origin_temp:YSW3 | -0.186 | 0.102 | 0.072 |
| origin_temp:YSW4 | -0.053 | 0.103 | 0.609 |
| origin_temp:YSW5 | -0.016 | 0.097 | 0.867 |
| origin_temp:YSW6 | -0.175 | 0.090 | 0.055 |
| origin_temp:YSW7 | 0.004 | 0.087 | 0.965 |
| origin_temp:YSW8 | -0.211 | 0.092 | 0.023 |
| origin_temp:YSW9 | -0.078 | 0.095 | 0.412 |
| origin_temp:YSW10 | 0.241 | 0.270 | 0.374 |
| origin_temp:YSW11 | -0.168 | 0.115 | 0.147 |
| origin_temp:YSW12 | -0.028 | 0.554 | 0.960 |
| origin_temp:YSW13 | -0.085 | 0.155 | 0.585 |
| origin_temp:YSW14 | 0.057 | 0.554 | 0.918 |
| origin_temp:YSW15 | NA | NA | NA |
| origin_temp:YSW16 | -0.184 | 0.263 | 0.485 |
| origin_temp:YSW17 | NA | NA | NA |

Supplementary Table 6.

| **Figure 1e, Model 6 - Response variable**: recovery lambda | | |  |
| --- | --- | --- | --- |
| **Generalized linear model with gamma distribution and log link**: (recovery growth rate) ~ (decline growth rate) * (early winter temperature) | | | |
| N=18 populations |  |  |  |
| *Term* | *Estimate* | *Std. error* | *P value* |
| Categorical temperature model – reference level: cold | |  |  |
| Intercept | 0.314 | 0.045 | <0.001 |
| gr_decline | -0.155 | 0.068 | 0.025 |
| tempcat-warm | -0.145 | 0.055 | 0.009 |
| gr_decline:tempcat-warm | 0.336 | 0.096 | <0.001 |
| Continuous temperature model | |  |  |
| Intercept | 0.381 | 0.129 | 0.004 |
| gr_decline | -0.314 | 0.221 | 0.157 |
| origin_temp | -0.023 | 0.019 | 0.232 |
| gr_decline:origin_temp | 0.049 | 0.033 | 0.140 |

Supplementary Table 7.

| **ED Figure 4, Model 7 - Response variable**: recovery lambda | | | |
| --- | --- | --- | --- |
| **Generalized linear model with gamma distribution and log link function**: (recovery growth rate) ~ (decline growth rate) | | | |
| N=8 populations | |  |  |
| *Term* | *Estimate* | *Std. error* | *P value* |
| Intercept | 0.284 | 0.016 | <0.001 |
| decline | -0.243 | 0.036 | <0.001 |

Supplementary Table 8.

| **ED Figure 5, Model 8 - Response variable**: daily pathogen growth rate | | | |
| --- | --- | --- | --- |
| **Generalized linear mixed model with gamma distribution and log link function**: (daily growth) ~ (origin) + (1\|cage_id) | | | |
| N=96 (24 experiment bats removed with no growth rate measurement) | | | |
| Reference level: MW1 | |  |  |
| *Term* | *Estimate* | *Std. error* | *P value* |
| Intercept | 0.069 | 0.007 | <0.001 |
| siteID:MW2 | -0.016 | 0.008 | 0.046 |
| siteID:NE1 | -0.024 | 0.009 | 0.007 |
| siteID:NE2 | -0.019 | 0.009 | 0.026 |
| siteID:NE3 | -0.019 | 0.009 | 0.042 |
| siteID:NE4 | -0.005 | 0.009 | 0.564 |
| siteID:NE5 | -0.008 | 0.009 | 0.382 |
| siteID:NE6 | -0.019 | 0.009 | 0.038 |

Supplemental Table 9.

| **Figure 2a&b, Model 9 - response variable**: daily pathogen growth rate | | | |
| --- | --- | --- | --- |
| **Generalized linear mixed model**: (daily pathogen growth) ~ log_10_(post-WNS minimum colony count) + (early winter origin site temperature) + (1\|cage_id), family=Gamma(link=log) | | | |
| N=96 (24 experiment bats removed with no growth rate measurement) | | | |
| *Term* | *Estimate* | *Std. error* | *P value* |
| Categorical temperature model – reference level: cold | | |  |
| Intercept | 0.043 | 0.009 | <0.001 |
| wnsminlog | 0.006 | 0.002 | 0.022 |
| tempcat:warm | -0.010 | 0.004 | 0.015 |
| Continuous temperature model | | |  |
| Intercept | 0.055 | 0.012 | <0.001 |
| wnsminlog | 0.007 | 0.003 | 0.007 |
| origin_temp | -0.003 | 0.002 | 0.044 |

Supplemental Table 10.

| **Figure 2c, Model 10 - response variable**: survival | | |  |
| --- | --- | --- | --- |
| **Generalized linear mixed model**: (survival) ~ (origin) + (1\|Cage_id), family=binomial (link=logit) | | | |
| Reference level: MW1 | | |  |
| N = 120 (all experimental bats) | | |  |
| *Term* | *Estimate* | *Std. error* | *P value* |
| Intercept | 1.008 | 0.830 | 0.225 |
| siteID-MW2 | -0.075 | 1.024 | 0.942 |
| siteID-NE1 | -1.402 | 1.032 | 0.174 |
| siteID-NE2 | -0.874 | 1.024 | 0.393 |
| siteID-NE3 | -0.904 | 1.028 | 0.379 |
| siteID-NE4 | 0.143 | 1.044 | 0.891 |
| siteID-NE5 | -2.175 | 1.094 | 0.047 |
| siteID-NE6 | -2.209 | 1.102 | 0.045 |

Supplemental Table 11.

| **Figure 2d, Model 11 - Response variable**: survival | |  |  |
| --- | --- | --- | --- |
| **Generalized linear mixed model**: (survival) ~ log10(pre-WNS colony count)*origin site early winter temp + (1\|Cage_id), family=binomial (link=logit) | | | |
| N=120 bats |  |  |  |
| *Term* | *Estimate* | *Std. error* | *P value* |
| CATEGORICAL MODEL |  |  |  |
| Intercept | -9.657 | 3.313 | 0.004 |
| prepoplog | 2.251 | 0.778 | 0.004 |
| tempcatwarm | 8.268 | 3.717 | 0.026 |
| prepoplog:tempcatwarm | -1.890 | 0.866 | 0.029 |
| CONTINUOUS MODEL |  |  |  |
| Intercept | -13.937 | 9.233 | 0.131 |
| prepoplog | 2.941 | 2.213 | 0.184 |
| origin_temp | 1.625 | 1.354 | 0.230 |
| prepoplog:origin_temp | -0.328 | 0.324 | 0.311 |

Supplemental Table 12.

| **ED Figure 6a, Model 12 - Response variable**: survival | |  |  |
| --- | --- | --- | --- |
| **Binomial generalized linear mixed model with logit link**: (survival) ~ (translocation site) + (1\|cage_id) | | | |
| N=120 bats | | |  |
| *Term* | *Estimate* | *Std. error* | *P value* |
| Intercept | 0.932 | 0.510 | 0.068 |
| Trans_site – NE-T | -1.135 | 0.584 | 0.052 |

Supplemental Table 13.

| **ED Figure 6b, Model 13 - Response variable**: survival | |  |  |
| --- | --- | --- | --- |
| **Binomial generalized linear mixed model with logit link**: (survival) ~ (early winter temperature) + Log_10_(early hibernation fungal load) + (early hibernation body mass) + (1\|cage_id) | | | |
| N=96 (experimental bats that entered Pd positive) | | |  |
| *Term* | *Estimate* | *Std. error* | *P value* |
| Categorical temperature model – reference level: cold | |  |  |
| Intercept | -12.938 | 3.530 | <0.001 |
| LOAD10 | -0.557 | 0.277 | 0.044 |
| mass | 1.051 | 0.345 | 0.002 |
| tempcat:cold | -0.793 | 0.488 | 0.104 |
| Continuous temperature model | | | |
| Intercept | -13.858 | 3.659 | <0.001 |
| LOAD10 | -0.539 | 0.273 | 0.048 |
| mass | 0.987 | 0.346 | 0.004 |
| origin_temp | 0.325 | 0.176 | 0.064 |

Supplemental Table 14.

| **Figure 3a&b, Model 14 - Response variable**: Log_10_ late hibernation fungal load | | | |
| --- | --- | --- | --- |
| **Generalized linear model**: (lgdL) ~ (WNS phase) * (early winter hibernation temperature) | | | |
| N=977 bats (historic n = 872, contemporary = 105) | |  |  |
| *Term* | *Estimate* | *Std. error* | *P value* |
| Categorical temperature model - reference level: phase-historic:tempcat-cold | | | |
| Intercept | -2.431 | 0.076 | <0.001 |
| phase-contemporary | 0.163 | 0.183 | 0.374 |
| tempcat-warm | 0.678 | 0.088 | <0.001 |
| phase-contemporary:tempcat-warm | -0.947 | 0.238 | <0.001 |
| Continuous temperature model – reference level: phase-historic | | | |
| Intercept | -3.673 | 0.157 | <0.001 |
| Phase-contemporary | 0.955 | 0.523 | 0.068 |
| Origin_temp | 0.236 | 0.020 | <0.001 |
| Phase-contemporary:origin_temp | -0.189 | 0.080 | 0.019 |

Supplementary Table 15.

| **Figure 3e&f, Model 15 - Response variable**: Log_10_ late hibernation fungal load | | | |
| --- | --- | --- | --- |
| **Generalized linear model**: (lgdL) ~ (status) * (early winter site temperature) | | | |
| N = 167 bats (free-flying n = 872, caged n = 105) | | |  |
| *Term* | *Estimate* | *Std. error* | *P value* |
| Categorical temperature model - reference level: status-caged:tempcat-cold | | | |
| Intercept | -1.810 | 0.176 | <0.001 |
| status-free | -0.458 | 0.222 | 0.040 |
| tempcat-warm | -0.452 | 0.234 | 0.053 |
| status-free:tempcat-warm | 0.183 | 0.295 | 0.535 |
| Continuous temperature model – reference level: status-caged | | | |
| Intercept | -0.328 | 0.634 | 0.605 |
| Status-free | -2.390 | 0.760 | 0.002 |
| Origin_temp | -0.266 | 0.095 | 0.005 |
| Status-free:origin_temp | 0.312 | 0.115 | 0.007 |

Supplementary Table 16.

| **ED Figure 8, Model 16 - Response variable**: Proportion of infected wing tissue | | | |
| --- | --- | --- | --- |
| **Generalized linear model**: (uv_photo) ~ (status) * (early winter site temperature) | | | |
| N = 167 bats (free-flying n = 872, caged n = 105) | | |  |
| *Term* | *Estimate* | *Std. error* | *P value* |
| Categorical temperature model - reference level: status-caged:tempcat-cold | | | |
| Intercept | -1.981 | 0.177 | <0.001 |
| status-free | -0.532 | 0.217 | 0.014 |
| tempcat-warm | 0.102 | 0.239 | 0.669 |
| status-free:tempcat-warm | -1.092 | 0.323 | <0.001 |
| Continuous temperature model – reference level: status-caged | | | |
| Intercept | -2.161 | 0.817 | 0.008 |
| Status-free | 2.404 | 0.836 | 0.004 |
| Origin_temp | 0.039 | 0.129 | 0.763 |
| Status-free:origin_temp | -0.417 | 0.136 | 0.002 |

**Supplementary Table 17. Summary table of figures, statistical analyses, and the corresponding results throughout the manuscript.** Figures and models are listed in the table in the order that they appear in the manuscript and extended data. Data types, sample sizes, and collection periods for each analysis can be found in the associated supplemental table as well as throughout the methods. Abbreviations included in the table are as follows: WNS = white-nose syndrome, ED = extended data, YSW = years since WNS arrival; epidemic period = years 1–4 following pathogen arrival, recovery period = years 8–17 following pathogen arrival; temperature = early winter origin site temperature (°C); WNS phase = historic (1-4 YSW) or contemporary (8–16 YSW); status = caged or free-flying bats. All models that explored the effect of early winter hibernation temperature were ran temperature as a continuous as well as categorical predictor variable (see supplemental tables for model output). NA values for distributions and variables indicate that the associated figure depicts raw data.

| **Figure** | **Model #** | **Table #** | **Distribution** | **Response Variable** | **Fixed effect** | **Random effect** | **Findings** |
| --- | --- | --- | --- | --- | --- | --- | --- |
| Fig. 1a | NA | NA | NA | NA | NA | NA | Mean colony decline was 92% for 17 of 18 sites (1 site has experienced no decline since WNS arrival); recovery ranges from 2.4–73.1% of pre-epidemic colony size |
| ED Fig. 1 | NA | NA | NA | NA | NA | NA | Raw colony count data across YSW for populations of different size categories |
| Fig. 1b | 1 | S1 | Gamma | Percent of pre-WNS population remaining during the epidemic | Temperature | NA | Warm sites experienced greater declines on average compared to cold sites |
| ED Fig. 2a | 2 | S2 | Poisson | Years to WNS minimum colony count | Temperature * log_10_ pre-WNS colony count | NA | Warm sites reached their minimum colony size prior to cold sites across all colony sizes |
| ED Fig. 2b | 3 | S3 | Gamma | Proportion of pre-WNS count at the minimum | Temperature * log_10_ pre-WNS colony count | NA | Warm sites experienced more severe declines than cold sites regardless of pre-epidemic colony size |
| Fig. 1c | 4 | S4 | Negative binomial | Percent difference between the epidemic and recovery periods | Temperature * log_10_ pre-WNS colony count | NA | Large warm sites have gained a larger percentage of their pre-WNS colony sizes during the recovery period (years 8–17) |
| Fig. 1d | 5 | S5 | Gamma | Annual population growth rate (𝜆) | Temperature * YSW[categorical] | NA | Warm sites stabilized after reaching their minimum colony size during the initial epidemic and are now consistently growing |
| Fig. 1e | 6 | S6 | Gamma | Recovery population growth rate (𝜆) | Decline population growth rate (𝜆) * Temperature | NA | The severity of initial declines interacts with environmental conditions to drive variable recovery |
| ED Fig. 3 | NA | NA | NA | NA | NA | NA | Temperature variation at origin and translocation sites over the hibernation season |
| ED Fig. 4 | 7 | S7 | Gamma | Recovery population growth rate (𝜆) | Decline population growth rate (𝜆) | NA | Decline and recovery population growth rates of *M. lucifugus* at translocation origin sites; populations that experienced more severe declines are now growing at greater rates |
| Fig. 2a, b | 9 | S9 | Gamma | Daily pathogen growth rate | Temperature * log_10_ post-WNS minimum colony count | Cage ID | Bats from populations with greater declines now experience reduced pathogen growth; bats from warm sites had lower pathogen growth on average |
| ED Fig. 5 | 8 | S8 | Gamma | Daily pathogen growth rate | Origin site | Cage ID | On-host pathogen growth rates varied by origin site; 10% of individuals reduced pathogen growth over the hibernation season |
| Fig. 2c | 10 | S10 | Binomial | Survival | Origin site | Cage ID | Overall survival was 51% and varied by origin site, ranging from 27-73% |
| Fig. 2d | 11 | S11 | Binomial | Survival | Temperature * log_10_ pre-WNS colony count | Cage ID | Individuals from warm sites had greater survival; pre-WNS colony count influenced survival for bats from cold sites |
| ED Fig. 6a | 12 | 12 | Binomial | Survival | Translocation site | Cage ID | Bats at the Midwest translocation site had higher survival due to less experimental time, although the difference was not significant |
| ED Fig. 6b | 13 | 13 | Binomial | Survival | Temperature + log_10_ early hibernation fungal load + early hibernation body mass | Cage ID | Bats originating from warm sites that entered the experiment with greater body mass and lower fungal loads had the highest survival |
| ED Fig. 7 | NA | NA | NA | NA | NA | NA | Early hibernation infection status and body condition across origin sites |
| Fig. 3a, b | 14 | 14 | Gaussian | log_10_ late hibernation fungal load | WNS phase * Temperature | NA | Late winter fungal loads of contemporary populations in warm sites are significantly lower compared to the epidemic period |
| Fig. 3c, d | NA | NA | NA | NA | NA | NA | Distribution of fungal loads from historic and contemporary populations |
| Fig. 3e, f | 15 | 15 | Gaussian | log_10_ late hibernation fungal load | Status * Temperature | NA | Bats originating from cold sites had the highest late winter fungal loads when translocated to warm sites, free-flying bats from warm sites had the lowest fungal loads |
| Fig. 3g | NA | NA | NA | NA | NA | NA | Temperature difference between cold and warm origin sites and the temperature of translocation sites |
| ED Fig. 8 | 16 | 16 | Gamma | Proportion of infected wing tissue | Status * Temperature | NA | Free-flying bats from cold sites had the highest degree of tissue invasion |
