## Extended Data Tables and Figures for "Environmental conditions drive selection and recovery following disease-induced declines"

**Extended Data Table 1.** Summary of common garden experiment origin site details. Each origin and translocation hibernacula were given an anonymized site code that reflects its geographic region within the United States (NE = Northeast, MW = Midwest). The number of bats represents the number of individual *Myotis lucifugus* collected from each origin site or moved to each translocation site (NE-T, MW-T). The mean early winter temperature (°C) was averaged from daily measurements (November–December). YOA = the estimated year of pathogen arrival at each site. The pre-WNS maximum colony count represents the most recent population census prior to pathogen arrival at each hibernaculum, and the post-WNS minimum colony count represents the lowest documented population colony size following pathogen arrival.

| Site Code | Number of bats | Mean early winter temperature (°C) | Year of pathogen arrival (YOA) | Pre-WNS maximum colony count | Post-WNS minimum colony count |
| --- | --- | --- | --- | --- | --- |
| NE1 | 15 | 7.48 | 2009 | 1,517 | 170 |
| NE2 | 15 | 6.13 | 2008 | 183,542 | 2,049 |
| NE3 | 15 | 7.74 | 2007 | 15,374 | 1,000 |
| NE4 | 15 | 5.62 | 2008 | 87,401 | 13,251 |
| NE5 | 15 | 4.58 | 2008 | 6,268 | 1,398 |
| NE6 | 15 | 4.55 | 2008 | 9,432 | 24 |
| NE-T | 90 | 7.58 |  | 835 | 0 |
| MW1 | 12 | 5.91 | 2015 | 20,101 | 23,516 |
| MW2 | 18 | 8.03 | 2015 | 143,000 | 2,747 |
| MW-T | 30 | 8.17 | 2015 | 241 | 0 |

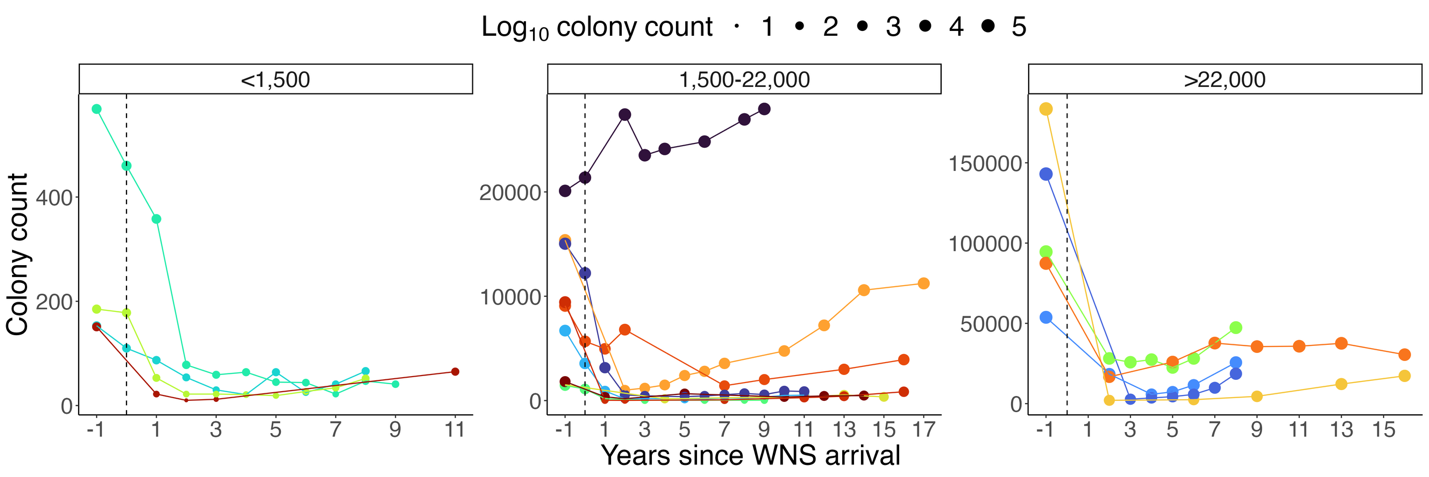

**Extended Data Figure 1**. Colony counts for *Myotis lucifugus* over years since white-nose syndrome (WNS) arrival. Population data were obtained from 18 hibernacula across the Northeast and Midwest US. Point size depicts the log_10_ colony count each year. The dashed vertical lines represent the year of WNS arrival. Sites are divided into size categories representing the pre-WNS maximum colony count (small: < 1,500; medium: 1,500_­_–22,000; large: > 22,000).

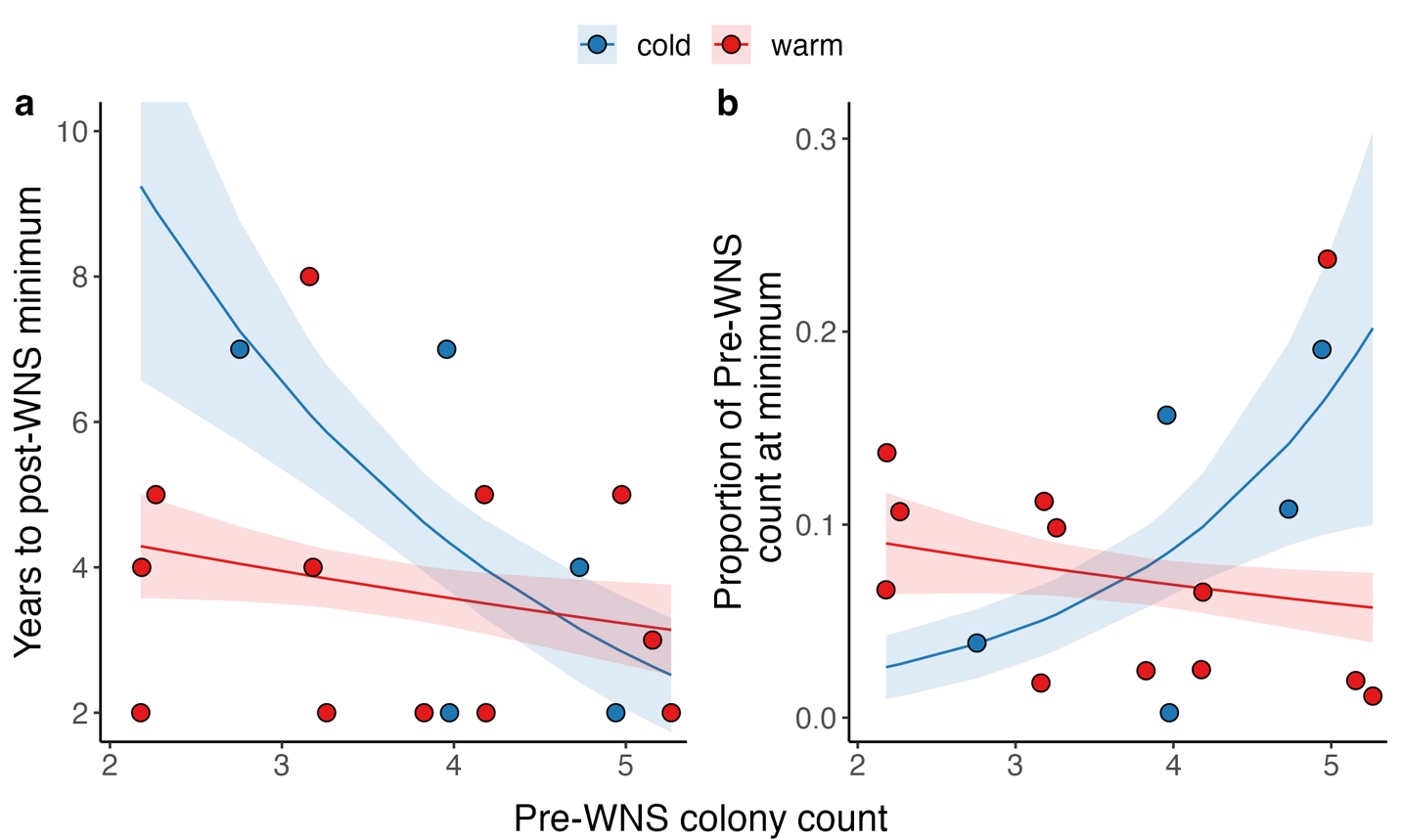

**Extended Data Figure 2**. Initial white-nose syndrome decline severity for little brown bat (*Myotis lucifugus*) populations. Relationship between early winter hibernation temperature, pre-WNS population size, and the a) years to reach post-WNS minimum colony count and b) proportion of pre-WNS population size at the minimum for 17 populations of *M. lucifugus* across the Northeast and Midwest US. Point color depicts the early winter site temperature (blue=cold, <6°C; red=warm, 6°C+ ). The transparent ribbons show the 95% confidence intervals. Model output can be found in Supplemental Tables 2 and 3.

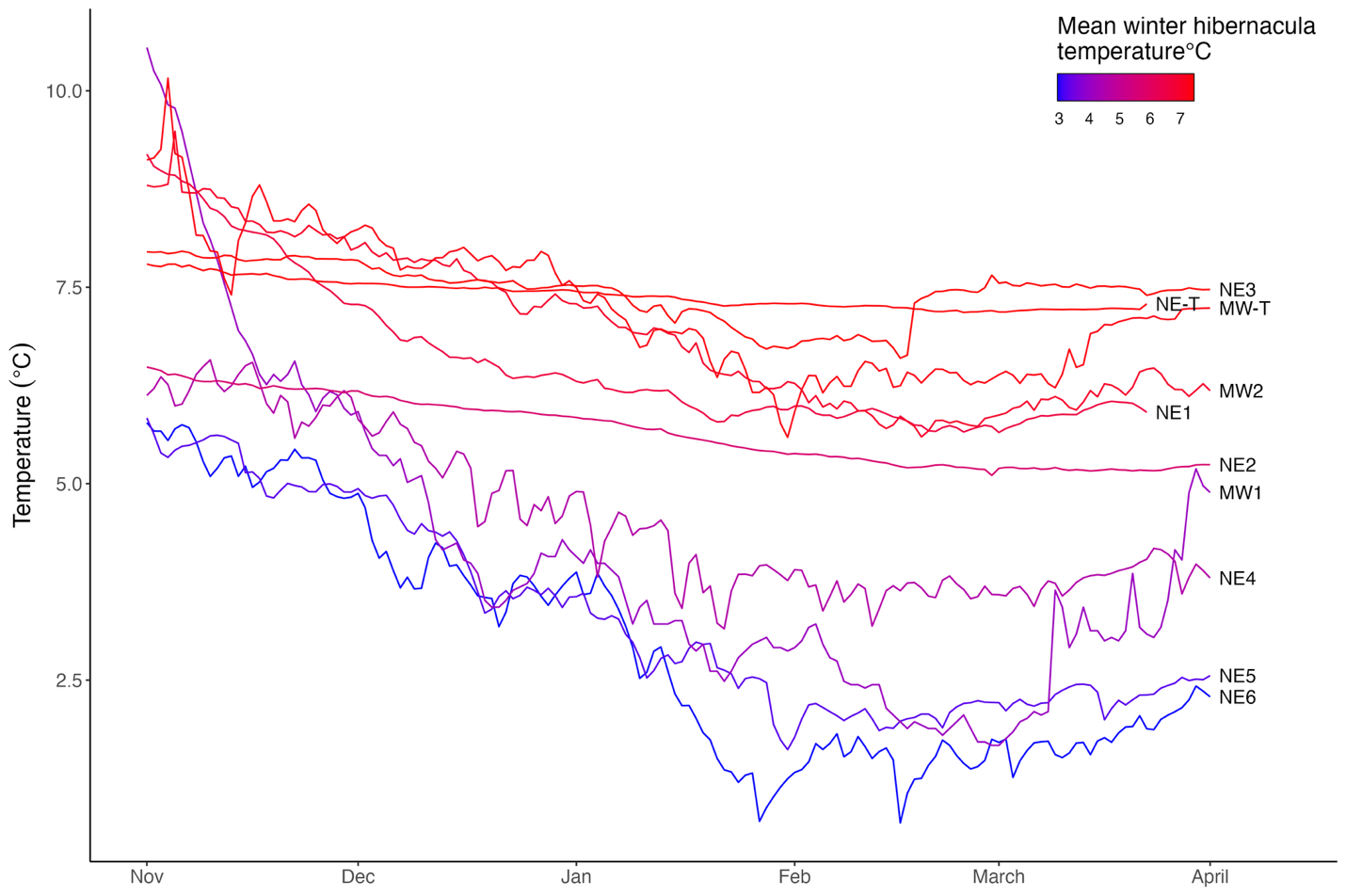

**Extended Data Figure 3**. Temperature variation at 8 origin and 2 translocation sites, plotted over time. The average winter temperature (Nov – March) in each site is as follows: NE1: 6.57 °C ± 0.95 SD, NE2: 5.67 °C ± 0.43 SD, NE3: 7.44 °C ± 0.35 SD, NE4: 4.57 °C ± 1.00 SD, NE5: 3.29 °C ± 1.24 SD, NE6: 2.95 °C ± 1.51, NE-T: 7.40 °C ± 0.18 SD, MW1: 4.11 °C ± 2.02 SD, MW2: 6.96 °C ± 0.99 SD, MW-T: 7.27 °C ± 0.88 SD.

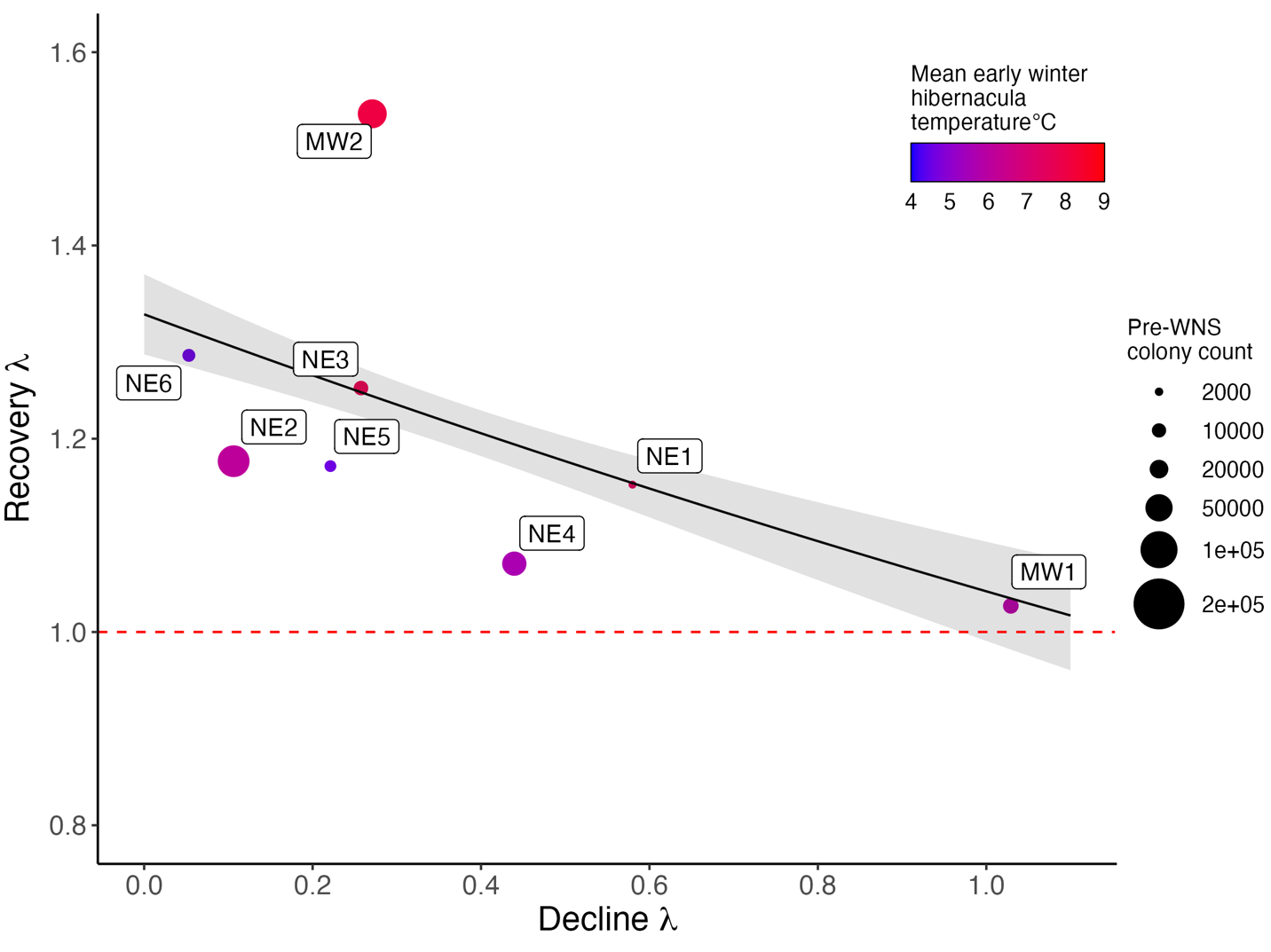
**Extended Data Figure 4**. Population growth rates (𝜆) of *M. lucifugus* at 8 origin sites during the initial decline vs. recovery periods. The recovery 𝜆 represents the population growth rate from the year of pathogen arrival till that site reached its minimum colony count (mean = 3.125 years following pathogen arrival, range = 2–7), while the decline 𝜆 represents the population growth rate from the minimum to the most recent count (range = 2–17 years following pathogen arrival). Point color represents the mean early winter hibernacula temperature for each origin site, and point size depicts the pre-WNS population size. The dashed red line indicates a stable population growth rate (𝜆=1). Populations that experienced more severe initial declines are now growing at greater rates (decline: -0.155 +/-0.036, p < 0.001, Supplemental Table 7).

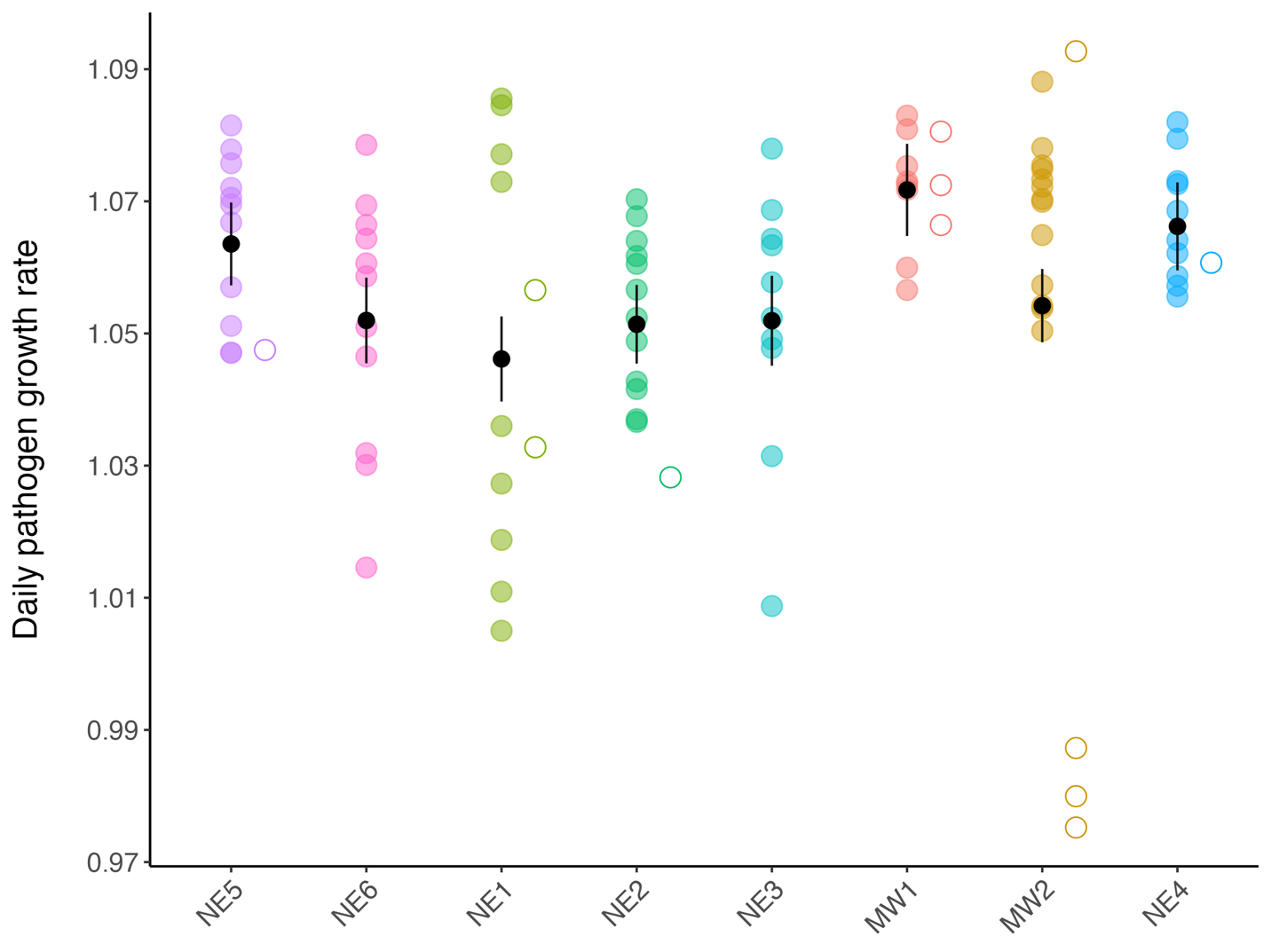

**Extended Data Figure 5.** Daily on-host pathogen growth rate by origin site. Sites are arranged on the x-axis according to survival (left=lowest, right=highest). Black points and error bars represent the model mean +/- standard error for each origin site (Supplemental Table 8). Open circles indicate individuals that reduced or cleared infections over the course of the experiment. Bats from site MW1 experienced the highest pathogen growth rates over the course of the experiment.

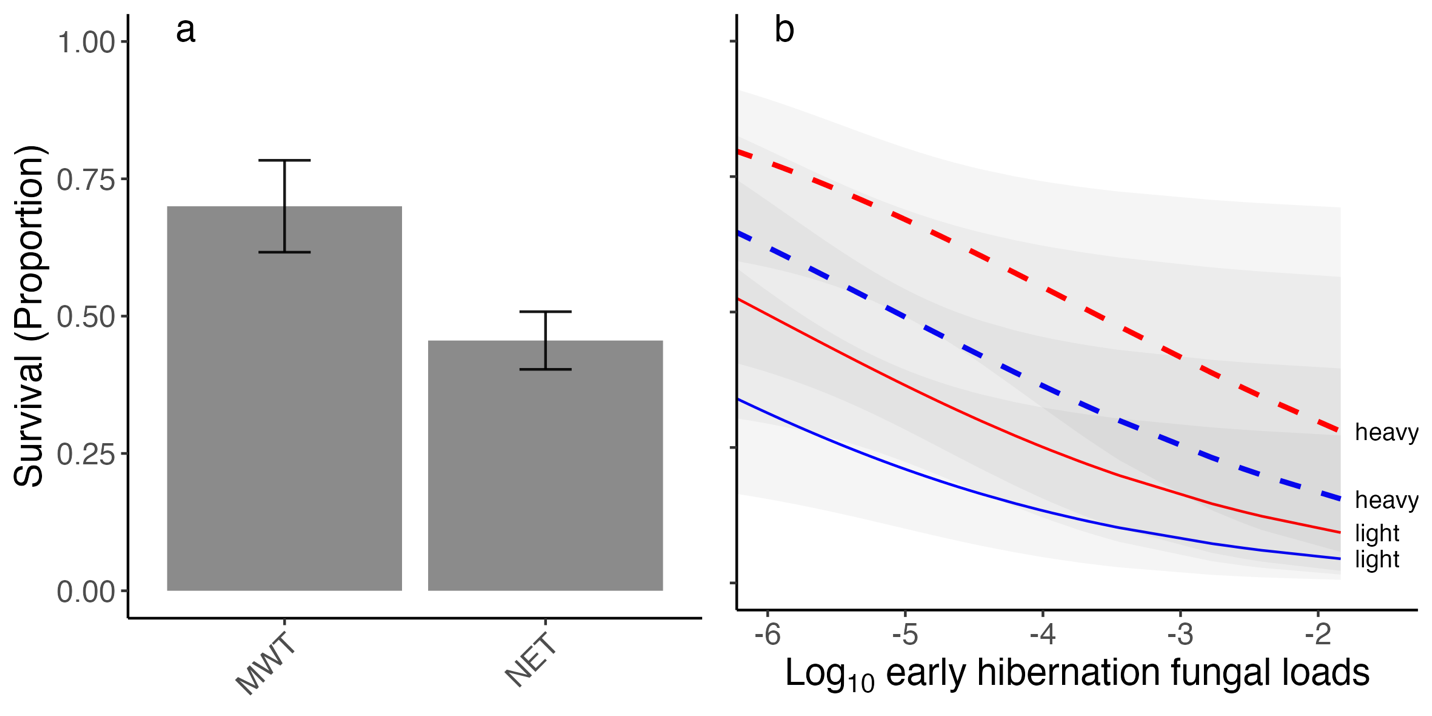

**Extended Data Figure 6**. Common garden experiment results. The relationship between survival and (a) translocation site and (b) origin site early winter temperature, early hibernation pathogen loads (ng of DNA) and early hibernation body mass. (a) Error bars represent the mean ± standard error for each translocation site (MWT = Midwest, NET = Northeast). (b) Lines depict the model prediction, and transparent ribbons show the 95% confidence intervals. Color of line represents whether bats originated from a cold (blue = <6°C) or warm (red = >6°C) hibernation site, and line type indicates model predictions for light (solid) or heavy (dotted) bats (individuals were divided into body mass categories based on the median weight at the onset of the experiment; median = 8.9 grams, range = 7.28–11.07 grams). Model output can be found in Supplemental Tables 12 and 13.

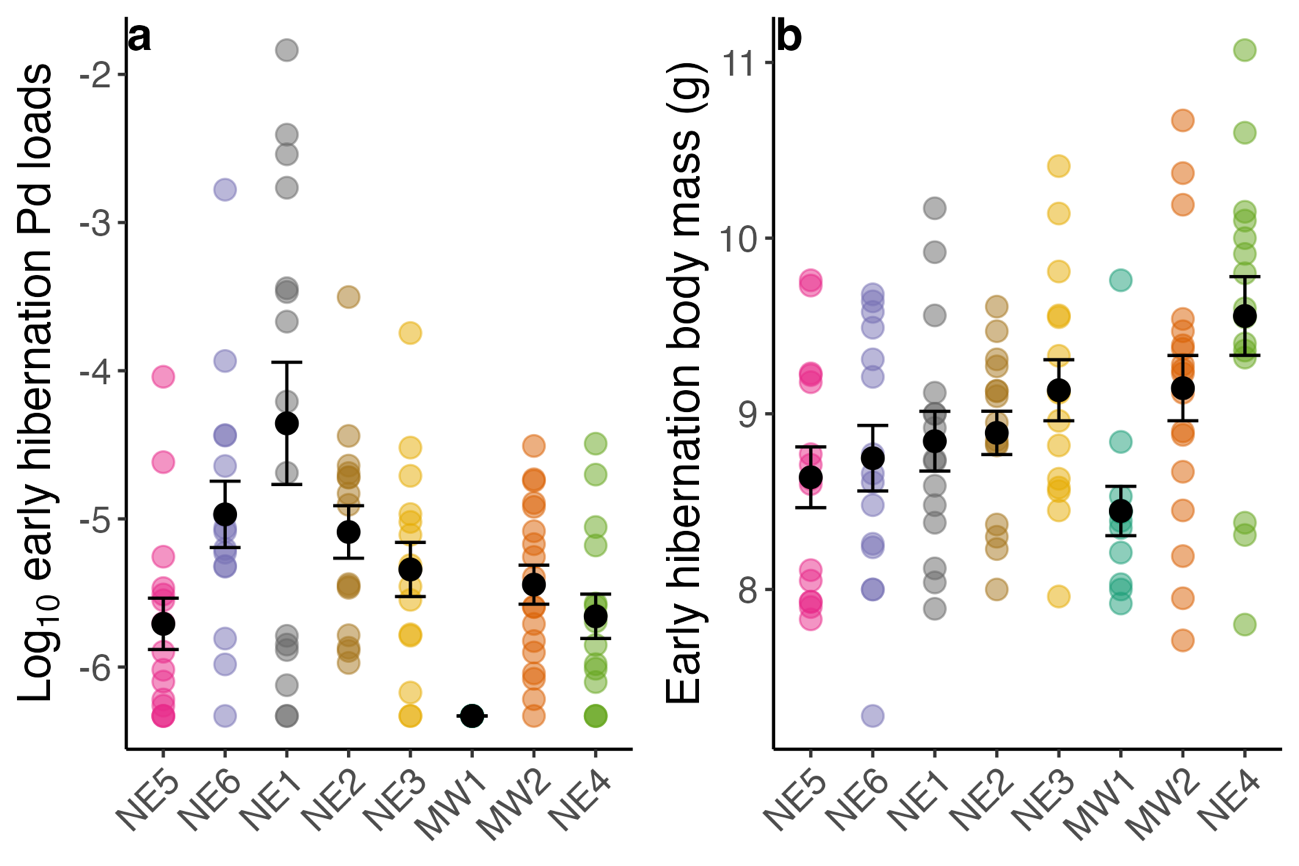

**Extended Data Figure 7**. Early hibernation infection status and body condition. (a) Pathogen (*Pseudogymnoascus destructans*, Pd) loads (ng of DNA) and (b) body mass (g) of bats at the onset of the common garden experiment. Sites are arranged on the x-axis according to survival (left=lowest, right=highest). Black points and error bars represent the mean +/- standard error for each origin site. Early hibernation on-host pathogen loads and body mass varied by origin site, and 24 (20%) individuals had no detectable fungus at the time of capture (a value of -6.33 represents a negative sample, equivalent to a C_t_ value of 40).

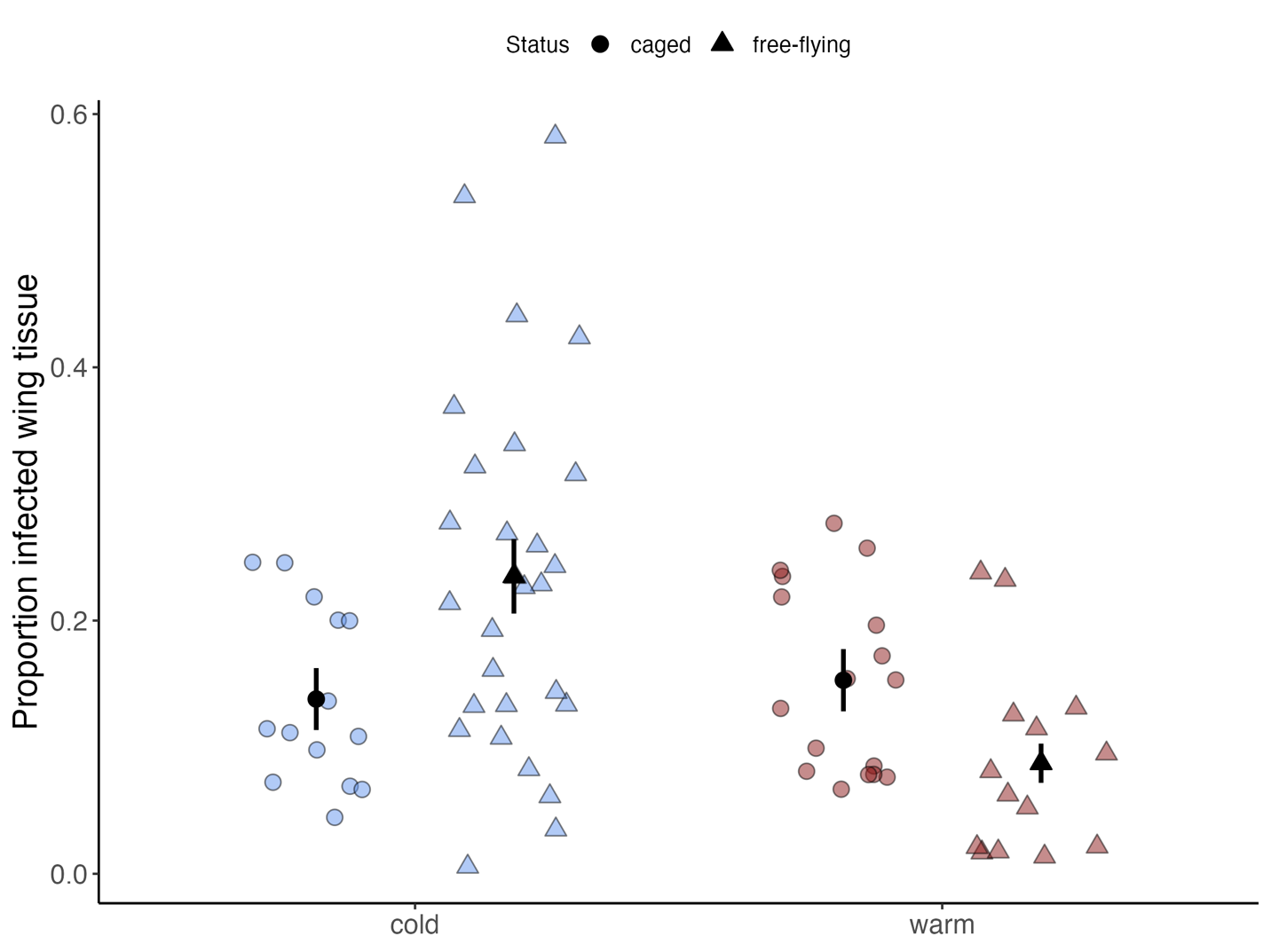

**Extended Data Figure 8.** Proportion of infected wing tissue as indicated by orange fluorescence on caged and free-flying bats in origin sites. Point colors represent the early winter temperature of origin sites (blue = cold, <6°C; red = warm, >6°C). Black points and error bats represent model estimates ± standard error (Supplemental Table 16). Point shape represents status (caged vs. free-flying). Free-flying bats were opportunistically captured at the end of hibernation at experiment termination within origin sites (n=7). Free-flying bats from cold sites had significantly higher levels of tissue invasion than free-flying bats from warm sites.
